# Structural insight into sodium ion pathway in the bacterial flagellar stator from marine *Vibrio*

**DOI:** 10.1101/2024.07.15.603494

**Authors:** Tatsuro Nishikino, Norihiro Takekawa, Jun-ichi Kishikawa, Mika Hirose, Seiji Kojima, Michio Homma, Takayuki Kato, Katsumi Imada

## Abstract

Many bacteria swim in liquid or swarm on surface using the flagellum rotated by a motor driven by specific ion flow. The motor consists of the rotor and stator, and the stator converts the energy of ion flow to mechanical rotation. However, the ion pathway and the mechanism of stator rotation coupled with specific ion flow are still obscure. Here, we determined the structures of the Na^+^-driven stator of *Vibrio*, namely PomAB, in the presence and absence of sodium ions and the structure with its specific inhibitor, phenamil, by cryo-electron microscopy. The structures and following functional analysis revealed the sodium ion pathway, the mechanism of ion selectivity, and the inhibition mechanism by phenamil. We propose a model of sodium ion flow coupled with the stator rotation based on the structures. This work provides insights into the molecular mechanisms of ion specificity and conversion of the electrochemical potential into mechanical functions.

## Introduction

Many motile bacteria convert electrochemical potential energy into cell motion using a filamentous organelle, the flagellum, which comprises the filament, the hook, and the motor (*1*). The flagellar filament is rotated by a motor embedded in the cell membrane at the base of the flagellum, and the filament rotation thrusts the cell, like a screw propeller. The motor can rotate both counterclockwise (CCW) and clockwise (CW) directions (*2*). The cell smoothly swims in CCW rotation and tumbles in CW rotation to change the swimming direction. The cell moves toward a favorable environment through the combination of the smooth swimming and tumbling modes.

The motor comprises the rotor and a dozen stators (*2*). The stator is an ion channel that converts the electrochemical potential of proton or sodium ions across the cell membrane. The rotor is composed of the MS-ring embedded in the cell membrane and the cytoplasmic C-ring attached below the MS-ring. The MS-ring is the assembly base for the flagellar structure, and the C-ring interacts with the stator to generate torque coupled with the specific ion flow through the stator.

The stator is a complex of two types of membrane proteins, the A- and B-subunits (*3*): MotA and MotB for the proton-driven stator (*4*), PomA and PomB for the sodium-driven stator from *Vibrio* species such as *Vibrio alginolyticus* (*5*), and MotP and MotS for the sodium-driven stator from some *Bacillus* species (*6*). The A-subunit (MotA/PomA/MotP) consists of four transmembrane helices and a cytoplasmic domain (*7*). The cytoplasmic domain contains highly conserved charged residues essential for torque generation and interacts with FliG, a C-ring component protein, to spin the rotor (*8–11*). The B-subunit (MotB/PomB/MotS) comprises a transmembrane helix and a C-terminal OmpA-like domain (*5*). The transmembrane helix contains a conserved aspartic acid residue essential for specific ion flux through the stator (*12*). The OmpA-like domain binds to the peptidoglycan layer or the T-ring to anchor the stator around the rotor (*13–15*). The transmembrane helix and the OmpA-like domain are connected by an amphiphilic helix, named “plug”, and a flexible linker. The plug helix controls open/close of the ion channel (*16–19*).

Recent structural studies using cryo-electron microscopy (cryoEM) have revealed that a stator complex consists of a pentameric A-subunit ring and a B-subunit dimer whose transmembrane helices are situated at the ring center (*20–22*). The pentameric ring can be formed by the A-subunit alone (*23*). Based on the stator structures, a model of the flagellar motor rotation has been proposed, in which the rotor is rotated by the A-subunit ring rotation around the B-subunit dimer driven by specific ion flux like a small gear turns a large gear wheel (*20–22, 24*). However, the ion pathway in the stator and the mechanism of stator rotation coupled with specific ion flux are still obscure.

The sodium-driven flagellar motor of *V. alginolyticus* (*Va*-motor) is a model system to study the molecular mechanism of torque generation coupled with specific ion flux because sodium ion is easier to detect, trace, and control than proton (*25*). The stator assembles to or disassembles from the *Va*-motor dependent on sodium concentration (*26*), and sodium-dependent structural change of the periplasmic region of PomB in the stator regulates the ion channel opening coupled with the stator anchoring around the rotor (*15, 27*) (**Fig. 1A**). Extensive genetic and biochemical studies of the *Va*-motor have identified the residues involved in ion conductivity, motor rotation, and interaction to the rotor. D24 of PomB directly binds sodium ions (*28*) and the D24N mutation impairs sodium ion flux (*29*), indicating that D24 is an essential residue for Na^+^ flow. T158 and T186 of PomA are also involved in the sodium ion conductivity (*30*). Recent cryoEM structure of the stator complex of *V. alginolyticus*, *Va*-PomAB, in 300 mM NaCl solution demonstrated that a sodium ion binds between T158 and T186 of PomA, which locate near D24, suggesting that the two threonine residues form a part of the sodium ion-selective filter (*22*). However, the proton-driven stator of *Rhodobacter sphaeroides* conserves the residues corresponding to D24, T158, and T186 (*5*), suggesting the presence of other factors for ion selectivity.

**Fig. 1.**
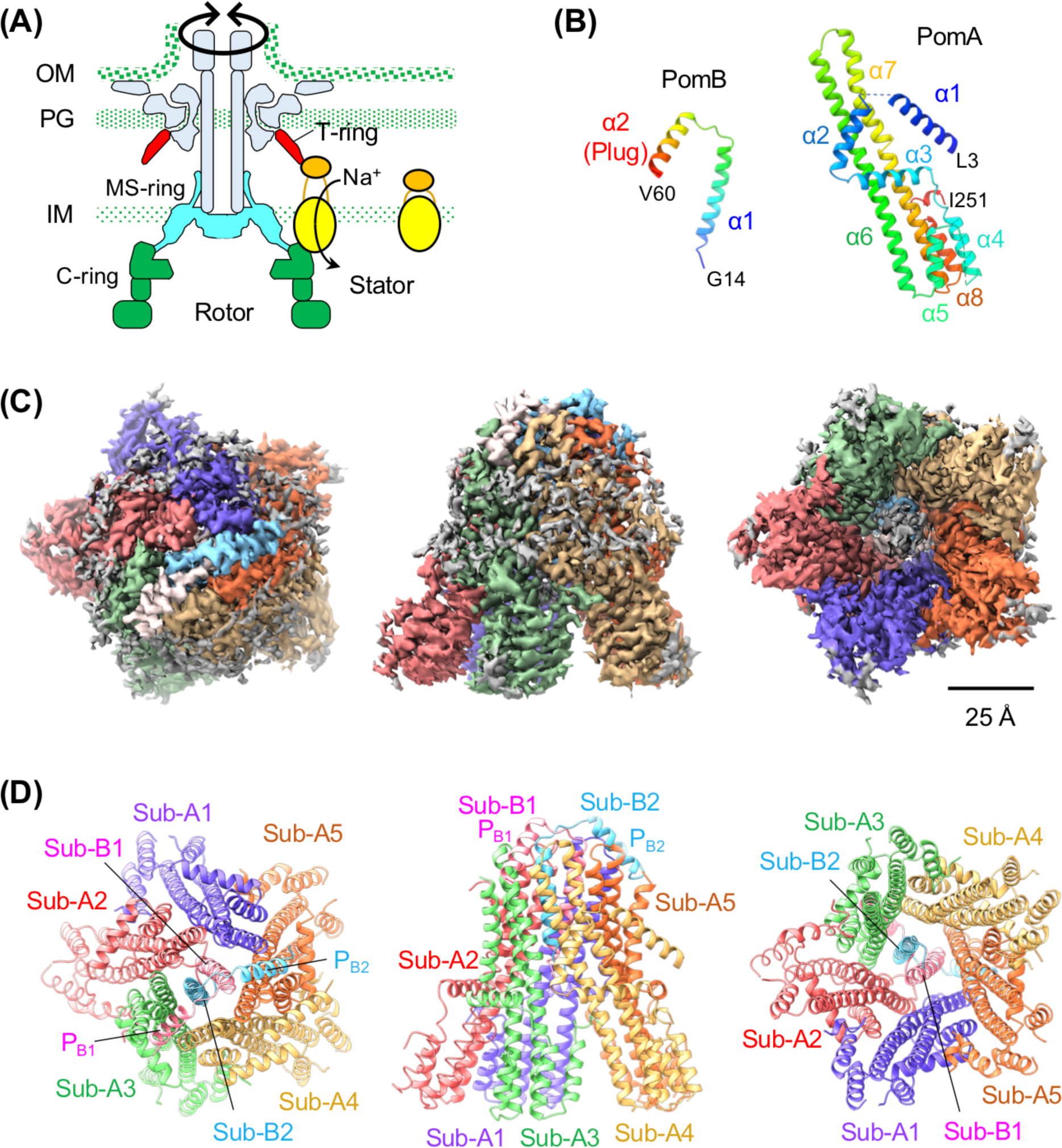
Structure of the stator of *V. alginolyticus*. **(A)** Schematic drawing of the polar flagellar motor of *V. alginolyticus*. The MS-, C-, and T-ring are colored in cyan, green, and red, respectively. Sodium ion changes the conformation of the periplasmic region of PomB (orange) to bind to the T-ring as well as to open the ion channel of the stator (yellow and orange). **(B)** Ribbon representation of PomA and PomB protomers. **(C)** CryoEM map of *Va*-PomAB viewed from periplasm (left), side (middle), and cytoplasm (right). PomB Sub-B1 and B2, and PomA Sub-A1, A2, A3, A4, and A5 are colored in light pink, sky blue, blue, red, light green, khaki, and orange, respectively. **(D)** Ribbon representation of *Va*-PomAB. The views and the subunit colors are the same as in **(C)**. The plug helices are labeled as PB1 and PB2.

To elucidate the sodium ion pathway, sodium ion selectivity, and the mechanism of specific ion flux coupled with the structural change of the stator, we determined the cryoEM structures of *Va*-PomAB and its PomB-D24N mutant stator (*Va*-PomAB(D24N)) in the presence and absence of sodium ion and compared the structures. Moreover, we also determined the structure of *Va*-PomAB in complex with phenamil, which is an amiloride analog and a specific inhibitor of sodium ion flow through *Va*-PomAB (*31*). We found two sodium ions in a tunnel along the cleft formed by two adjacent PomA subunits and therefore assigned the tunnel as the sodium ion pathway. The tunnel consists of the hydrophobic and hydrophilic cavities separated by a bottleneck region. We show that the hydrophobic cavity and the bottleneck region contribute to select sodium ions. Phenamil inhibits the sodium ion flow not by blocking the ion pathway but by disturbing the PomA rotation. We finally propose a model of sodium ion flux coupled with the PomA ring rotation.

## Results

### Structure of *Va*-PomAB

To investigate the sodium ion pathway, we determined the structures of *Va*-PomAB and *Va*-PomAB(D24N) using cryoEM. *Va*-PomAB and *Va*-PomAB(D24N) were expressed with a hexa-histidine tag at the C-terminus of PomB and purified by Ni affinity chromatography followed by size exclusion chromatography with lauryl maltose neopentyl glycol (LMNG) and 100 mM NaCl (fig. S1, A and B).

The cryoEM structure of *Va*-PomAB was determined at 3.12 Å resolution (**Fig. 1, B to D**). The transmembrane helices of the PomB dimer (Sub-B1 and B2) penetrate the central hole of the pentameric PomA ring (Sub-A1 to A5), as reported by Hu et al (*22*). The PomA protomer consists of eight α-helices (α1-8) (**Fig. 1B** and fig. S1C). α1 except for that in Sub-A4, the loops between α1 and α2 and between α4 and α5, and the C-terminal two residues were invisible in the cryoEM density map and therefore were not modeled (**Fig. 1, B to D** and fig. S1C). The PomB protomer was modeled from residues G14-V60, which include the transmembrane helix (α1) and the plug helix (α2) (**Fig. 1B** and fig. S1D). The N-terminal 13 residues and the C-terminal 255 residues (Q61-Q315) including the periplasmic domain were invisible in the density map (**Fig. 1, B to D**, and fig. S1D). The plug helix fits into the cleft between the periplasmic regions of two adjacent PomA protomers; Sub-A1 and A5 for Sub-B1, and Sub-A3 and A4 for Sub-B2 (**Fig. 1, C and D**). Due to the interaction, the periplasmic chains of α6 and α7 (G157-A182) in Sub-A3 and Sub-A5 are slightly twisted, although the root mean square deviation (RMSD) values for Cα atoms of any pair of PomA protomers are less than 1 Å. Therefore, PomA subunits are arranged in a slightly distorted pentagonal shape at the periplasmic side, whereas the remaining parts are arranged in a regular pentagon (**Fig. 1, C and D**). This is in contrast to the previous PomAB structure by Hu et al (*22*), in which the PomA forms an asymmetric pentameric ring (fig. S1E).

D24 in α1 of PomB is essential for the sodium ion flux and directly binds sodium ions (*28*). Since the α1 helices of the PomB dimer form a parallel coiled-coil structure, two sodium ion pathways are expected along the α1 helices of the PomB dimer, and we found a long and a short narrow tunnel including D24 of Sub-B1 and B2, respectively, in *Va*-PomAB (**Fig. 2**).

**Fig. 2.**
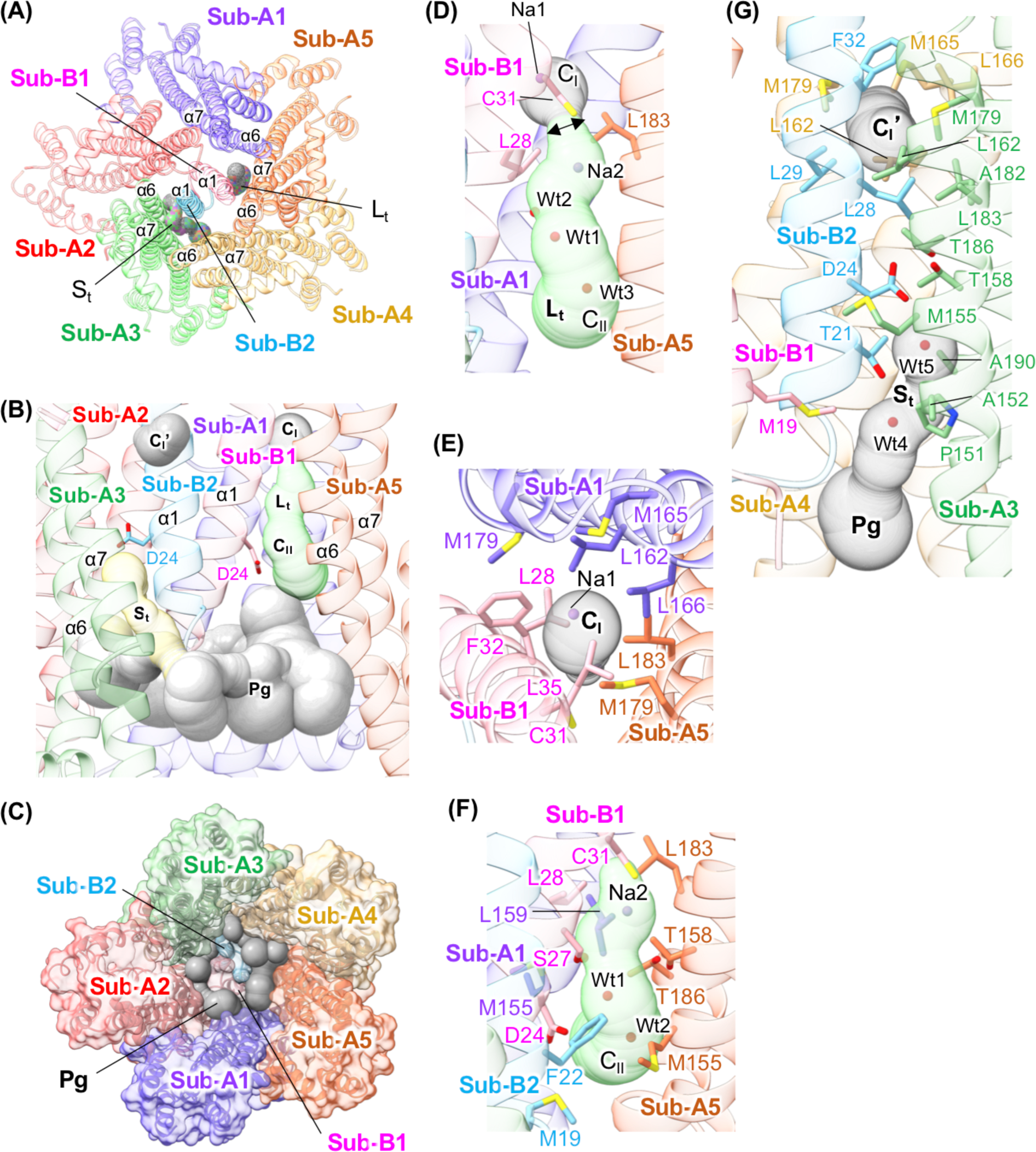
The long and short tunnels in *Va*-PomAB. The subunits are colored as in Fig. 1D. **(A)** Position of the long (Lt) and short (St) tunnels in *Va*-PomAB viewed from the periplasmic side. The residues from 40 to 60 including the plug helices are removed for easy viewing. The tunnels are shown as gray blobs. **(B)** The long (Lt) and short (St) tunnels are viewed from the bottom of **(A)**. Sub-A4 is removed for easy viewing. D24 residues of PomB are shown by stick model. Cavity-I (CI), Cavity-I’ (CI’), and the pentagonal groove (Pg) are shown by gray blobs. Cavity-II (CII) is indicated by green blobs. The short tunnel (St) is drawn by yellow blobs. **(C)** Cytoplasmic view of *Va*-PomAB. **(D)** Close-up side view of the long tunnel (Lt). The thin bottleneck region is indicated by a two-headed arrow. The residues involved in the bottleneck region are shown in the stick model. Sodium ions (Na1 and Na2) and solvent molecules (Wt1, Wt2, and Wt3) in the long tunnel are indicated by balls. **(E)** Close-up view of Cavity-I (CI) viewed from the periplasmic side. The residues that form the inner wall of Cavity-I are shown by the stick model. **(F)** Close-up side view of Cavity-II (CII). The residues that form the inner wall of Cavity-II are shown by the stick model. **(G)** Close-up side view of the short tunnel (St) and Cavity-I’ (CI’). The residues that form the inner wall of the short tunnel and Cavity-I’ are shown by stick model.

The long tunnel extends along the cleft composed of helices α6 and α7 of Sub-A1 and A5, and α1 of Sub-B1 covers the cleft to form the tunnel (**Fig. 2, A, B, and D**). The cytoplasmic side of the tunnel leads to the pentagonal groove open to the cytoplasm (**Fig. 2, B and C**), whereas the periplasmic side of the tunnel is closed (**Fig. 2D**). The tunnel is divided into two sections, Cavity-I in the periplasmic side and Cavity-II in the cytoplasmic side, by the thin bottle neck region formed by L28 and C31of Sub-B1 and L183 of Sub-A5 (**Fig. 2D**). Cavity-I is an ellipsoidal fully hydrophobic space with an approximate diameter of 6 Å (considering hydrogen atoms) composed of α1 of Sub-B1, α6 and α7 of Sub-A1, and α7 of Sub-A5 (**Fig. 2, B, D, and E**). The inner wall of Cavity-I is formed by M179 and L183 of Sub-A5, L162, M165, L166, and M179 of Sub-A1, and L28, C31, F32, and L35 of Sub-B1. Cavity-I contains a large solvent density at its center (**Fig. 3A**). In contrast to Cavity-I, Cavity-II is a hydrophilic space consisting of α1 of Sub-B1, α6 and α7 of Sub-A5, and α6 of Sub-A1 (**Fig. 2, B, D, and F**). T158 and T186 of Sub-A5, and S27 and D24 of Sub-B1 contribute to the hydrophilic nature of the cavity. L159 of Sub-A1, L183 of Sub-A5, L28 and C31 of Sub-B1, and F22 of Sub-B2 also form the inner wall of Cavity-II. G154 and M155 of Sub-A5, M155 of Sub-A1, D24 of Sub-B1, and M19 of Sub-B2 are located around the cytoplasmic entrance of Cavity-II. The side chain carboxy group of D24 points toward the cytoplasmic space. Cavity-II contains a few density peaks corresponding to solvent molecules or ions (**Fig. 3B**). Hu et al reported that a sodium ion is bound between T158 and T186 (*22*), but we found no extra density between the residues. This difference is probably because their structure was determined at 300 mM NaCl whereas our structure was at 100 mM NaCl.

**Fig. 3.**
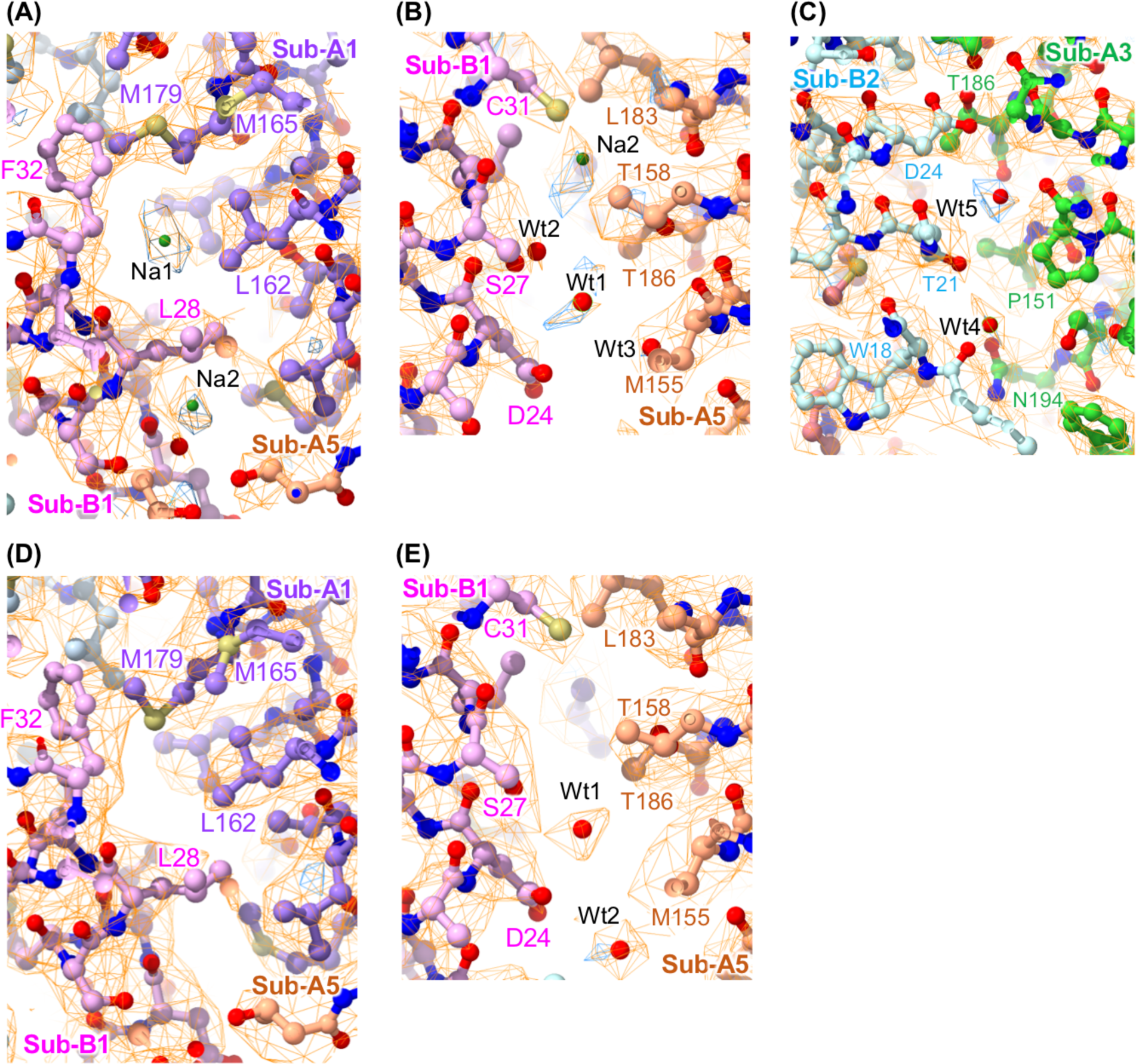
Ion and solvent molecules found in the tunnels. The cryoEM map (orange) and the difference map (cryo-EM map minus protein model map) (blue) are shown with the atomic model. Close-up view of **(A)** the long tunnel viewed from the periplasmic side, **(B)** Cavity-II, and **(C)** the short tunnel of *Va*-PomAB at 100 mM NaCl. **(D)** Close-up view around the long tunnel viewed from the periplasmic side and **(E)** Cavity-II of *Va*-PomAB at 100 mM KCl. Sodium ions (Na1 and Na2) and solvent molecules (Wt1 and Wt2) are indicated by balls

Unlike Sub-B1, D24 and L28 of Sub-B2 are closely in contact with α6 and α7 of Sub-A3. Therefore, only a short narrow tunnel to the cytoplasmic pentagonal groove exists below D24 of Sub-B2 (**Fig. 2, A, B, and G**). The short narrow tunnel formed by P151, A152, M155, G189, A190 of Sub-A3, and M19, G20, and T21 of Sub-B2. Two significant solvent density peaks exist in the tunnel (**Fig. 3C**). The sidechain of D24 of Sub-B2 is surrounded by M155, T158, and T186 of Sub-A3 and hydrogen bonds with T158 and T186. L28 of Sub-B2 hydrophobically interacts with L162, M179, A182, and L183 of Sub-A3. Above L28, there is an isolated hydrophobic cavity (Cavity-I’) with a similar size of Cavity-I at equivalent height to Cavity-I (**Fig. 2, B and G**). The Cavity-I’ is formed by M179 and L183 of Sub-A3, L162, M165, L166, and M179 of Sub-A4, and L28, L29, and F32 of Sub-B2. Because of the symmetry mismatch between PomA and PomB, the PomB residues that form Cavity-I’ differ from those of Cavity-I.

### Structure of *Va*-PomAB(D24N)

The cryoEM structure of *Va*-PomAB(D24N) was determined at 3.16 Å resolution (fig. S2B). The overall structure is very similar to that of *Va*-PomAB (root-mean-square-deviation value for corresponding Cα atoms: 0.3 Å). The side chain conformations of the residues in the two tunnels are almost identical to those of *Va*-PomAB except for N24 of Sub-B2 and M155 of Sub-A3. The side chain amide of N24 faces toward M155, and the side chain tip of M155 turns to avoid steric hindrance to N24. The solvent densities in Cavity-I, Cavity-II, and the short narrow tunnel are also found in PomAB(D24N).

### Identification of sodium ion

To identify sodium ions in *Va*-PomAB, we determined the structure of *Va*-PomAB at 100 mM KCl at 3.43 Å resolution and compared it with the structure at 100 mM NaCl. No significant overall structural differences were observed in the presence and absence of sodium ions (**Fig. 1D** and fig. S2A). However, the two strong densities in Cavity-I and Cavity-II disappeared in the structure at 100 mM KCl (**Fig. 3, D and E**). We therefore assigned the two densities as sodium ions and the other solvent densities as water molecules. We also determined *Va*-PomAB(D24N) at 100 mM KCl at 3.42 Å and obtained similar results to wild-type *Va*-PomAB. The structure is almost identical to *Va*-PomAB(D24N) at 100 mM NaCl (fig. S2, B and C) but the two strong densities in Cavity-I and Cavity-II disappeared, supporting that the two strong densities correspond to sodium ion.

### Structure of *Va*-PomAB with phenamil

Since phenamil inhibits the polar flagellar motility of *Vibrio* by inhibition of sodium ion flux through *Va*-PomAB (*31–33*), it would bind to the sodium ion pathway. We, therefore, determined the structure of *Va*-PomAB with phenamil and 100 mM NaCl at 3.32 Å resolution. The PomA subunits of the phenamil-bound *Va*-PomAB are asymmetrically arranged on the cytoplasmic side (**Fig. 4A**). Compared with the structure of *Va*-PomAB without phenamil, the cytoplasmic domain of Sub-A1 moves toward inside of the ring, and that of Sub-A5 and A2 outward and slightly left (viewed from the bottom) of the ring (fig. S1E). This subunit arrangement resembles the PomAB structure by Hu et al (*22*) (fig. S1E), suggesting that this subunit arrangement is not induced by phenamil binding. On the other hand, no significant difference in the conformation of each corresponding PomA protomer with and without phenamil. The RMSD values for Cα atoms of each protomer are less than 0.73 Å (Sub-A1, 0.73 Å; Sub-A2, 0.66 Å; Sub-A3, 0.44 Å; Sub-A4, 0.39 Å; Sub-A5, 0.68 Å), indicating that phenamil binding does not induce a significant conformational change of PomA.

**Fig. 4.**
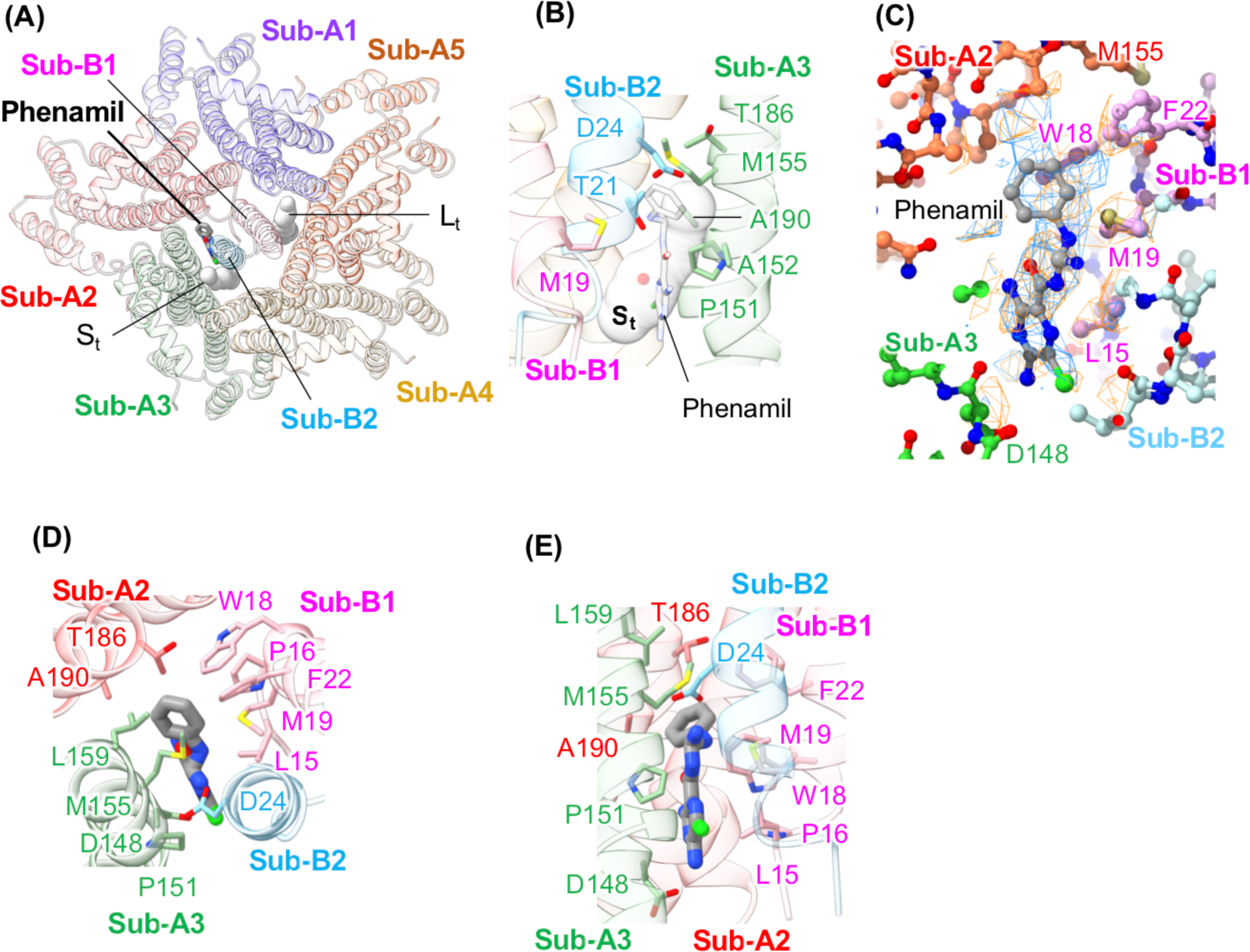
Structure of *Va*-PomAB with Phenamil. **(A)** Ribbon representation of *Va*-PomAB with phenamil viewed from the periplasmic side. The bound phenamil is shown by the stick model. The residues from 40 to 60 including the plug helices are removed for easy viewing. The long (Lt) and short narrow (St) tunnels are shown as gray blobs. **(B)** Phenamil binds near the short narrow tunnel. Phenamil and the residues that form the inner wall of the short tunnel are shown by the stick model. **(C)** The CryoEM map (orange) and the difference map (cryo-EM map minus protein model map) (blue) are shown with the atomic model. **(D and E)** Close-up view of the phenamil binding site from periplasmic view **(D)** and the side view **(E)**. Phenamil and the residues that interact with phenamil are shown by the stick model. The carbon atoms of phenamil are colored gray. Oxygen, nitrogen, chloride, and hydrogen atoms are colored red, blue, green, and white, respectively. The subunits are colored as in Fig. 1D.

An elongated density corresponding to phenamil appeared near the short narrow tunnel just below D24 of Sub-B2 (**Fig. 4, A to C**). The density is located in the cleft formed by Sub-B1, B2, and A3. Although the density is rather unclear, we were able to fit a phenamil model in the density (**Fig. 4C**). The phenyl group of phenamil is accommodated in the hydrophobic pocket formed by W18 and F22 of Sub-B1, T186 and A190 of Sub-A2, and M155 and L159 of Sub-A3 (**Fig. 4, D and E**). The imino group interacts with D24 of Sub-B2, whose side chain conformation is similar to D24N. The chloro pyrazinecarboxamide blocks the short narrow tunnel and the chlorine gets stuck in the tunnel (**Fig. 4, B and E**). The pyrazine ring forms a CH-pi interaction with L15 of Sub-B1. The amino group at position 5 of the pyrazine ring is located within a distance that can electrostatically interact with D148 of Sub-A3.

No extra density corresponding to phenamil was observed around the long tunnel, although the two sodium densities were observed in Cavity-I and Cavity-II. Since phenamil binds in the gap between PomA and PomB near D24 of Sub-B2, which is in the fully closed one of the two possible sodium pathways, phenamil does not directly block the sodium ion pathway but may disturb the PomA rotation coupled with sodium ion flux.

### Sodium ion pathway

The previous studies showed that D24 of PomB binds sodium ions (*28*), and D24N mutation completely inhibited sodium flux (*29*). Therefore, D24 should be located in the sodium ion pathway. D24 of Sub-B1 is in the long tunnel containing sodium ions in Cavity-I and Cavity-II (**Fig. 2**). T158 and T186, which are also essential for sodium flux (*30*), are located on Cavity-II. Thus, the long tunnel along Sub-B1 is the most plausible candidate for the sodium ion pathway. However, the tunnel has no periplasmic entrance for sodium ions, even if the plug region is removed from the model. Therefore, some structural change upon plug opening is needed for the sodium ion uptake. The plug regions slightly distort the periplasmic sides of α6 and α7 in Sub-A3 and A5. Thus, the plug opening may induce the structural change of the helices for sodium ion uptake.

### Cavity-I is a size filter for sodium ions

Since Cavity-I is fully hydrophobic, the sodium ion would be hydrated in Cavity-I. Previous experimental and simulation studies showed that the first hydrated shell peak position of Na^+^-O is 2.30 - 2.39 Å and that of Na^+^-H is 3.0 Å (*34*). The diameter of Cavity-I is about 6 Å (considering the size of C-H). Thus, Cavity-I can accommodate a hydrated sodium ion. In contrast, the first hydrated shell peak position of K^+^-O is 2.7-2.8 Å and that of K^+^-H is 3.4 Å (*34*). The hydrated potassium ion is about 1 Å larger than the hydrated sodium ion in diameter and is larger than Cavity-I. Thus, potassium ions cannot get into Cavity-I. Similarly, a hydrated oxonium ion (eigen cation or H3O^+^(H2O)3) (*35*) may be impossible to go into Cavity-I, because the O-O distance between H3O^+^ and H2O hydrating the eigen cation (*36*) is 2.58 - 2.60 Å, and thus a hydrated oxonium ion is 0.5 Å larger than a hydrated sodium ion. On the other hand, the Li^+^-O distance of hydrated lithium ion (*37*) is 1.90 - 2.17 Å, suggesting that the hydrated lithium ion can be accommodated in Cavity-I. The *Vibrio* motor can be driven by lithium ion (*38*), but not by potassium ion or oxonium ion. These facts suggest that Cavity-I is not a simple ion path but contributes to the selection of sodium ions as a size filter.

To examine this idea, we introduced mutations in L166 and M179 of PomA in the plug deletion stator and observed the growth of the mutant cells in the presence of NaCl to evaluate the sodium ion conductivity (**Fig. 5**). L166 and two M179 residues in the adjacent PomA subunits are located on the cleft that forms the middle wall of Cavity-I. No growth inhibition was observed in L166K and M179K mutant cells, indicating that these mutations completely block the sodium ion flux. L166A and M179A mutant cells showed growth inhibition but less than the plug deletion mutant cells. These results suggest that Cavity-I is a part of the sodium ion pathway and its size greatly affects that sodium ion flux. A previous study showed that M179C mutation in PomA conferred a non-motile phenotype (*39*). Moreover, M179 is conserved in the Na^+^-type stator, while the corresponding residue of the H^+^-type stator is isoleucine (**Fig. 5C**). These previous results are consistent with our results.

**Fig. 5.**
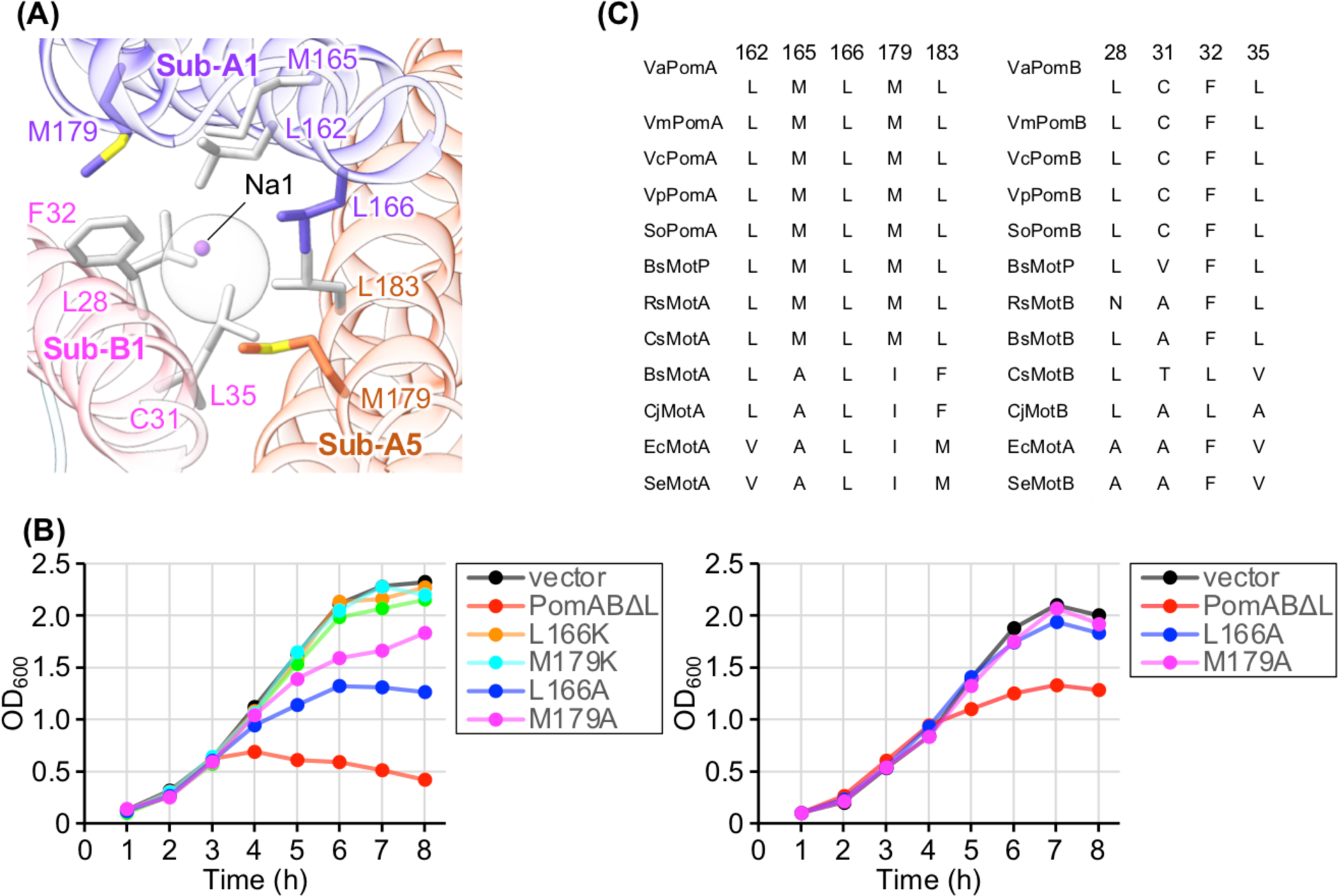
The Na^+^ conductivity of the stators with mutation at PomA L166 or M179. **(A)** Close-up view of Cavity I in *Va*-PomAB. The residues on the surface of Cavity I are labeled. The subunits are colored as in Fig. 1D. **(B)** Effects of PomA mutations on cell growth by plug deletion stators. *E. coli* cells harboring the plasmids pBAD33 (vector) or pTSK37 (PomABΔL or with various mutations) were cultured in LB medium with the final Na concentration of 102 mM (left panel) or LB-Na0 medium with the final Na concentration of 17 mM (right panel) at 37° C for 7 h. **(C)** Comparison of amino acids on the surface of Cavity-I. The sequences are *Vibrio alginolyticus*, VaPomA and B; *Vibrio mimicus,* VmPomA and B; *Vibrio*. cholerae, VcPomA and B; *Vibrio*. *parahaemolyticus,* VpPomA and B; *Shewanella oneidensis*, SoPomA and B; *Bacillus subtilis*, BsMotP and S, or MotA and B; *Rhodobacter sphaeroides*, RsPomA and B; *Campylobacter jejuni*, CjMotA and B; *Escherichia coli*, EcMotA and B; *Salmonella enterica*, SeMotA and B.

## Discussion

Recent structural studies on the stator have revealed that the A5B2 architecture is conserved (*20–22*). Based on the structures, a stator rotation model has been proposed for the torque generation mechanism of the flagellar motor (*20–22*). In the model, the stator A-subunit ring rotates around the PomB dimer in the CW direction viewed from the periplasmic side, and the rotor is rotated by interacting with the cytoplasmic domains of the A-subunit ring. Since the flagellar motor is driven by ion motive force, the A-subunit ring rotation should be coupled with ion flux through the stator complex. In this study, we determined the Na^+^-driven PomAB stator structures and identified sodium ions and the sodium ion pathway. Based on the structures, we propose a model of sodium ion flux coupled with PomA rotation (**Fig. 6**). We found two sodium ions in the long tunnel composed of α1 of Sub-B1 and the cleft formed by two adjacent PomA subunits, Sub-A1 and A5. Since PomA forms a pentamer ring in the stator, five equivalent PomA clefts exist in a stator complex. They interact with different surfaces of PomB, and therefore, are in different states (a, b, c, d, and e in **Fig. 6**). The cleft at state a forms the long tunnel including Cavity-I and -II, and that at state d forms the short narrow tunnel and Cavity-I’. The space corresponding to Cavity-I at state b is filled by L20 of Sub-B1, at state c by C31 of Sub-B2, and at state e by M30 of Sub-B1 and F33 of Sub-B2. Thus, the clefts at state b, c, and e have no cavity corresponding to Cavity-I. According to the stator rotation model, the state of each cleft changes with PomA rotation. For example, the cleft formed by Sub-A3 and A4 (the A3A4-cleft) transitions its state d, a, c, e, and b with every 36 degrees rotation in this order (**Fig. 6**).

**Fig. 6.**
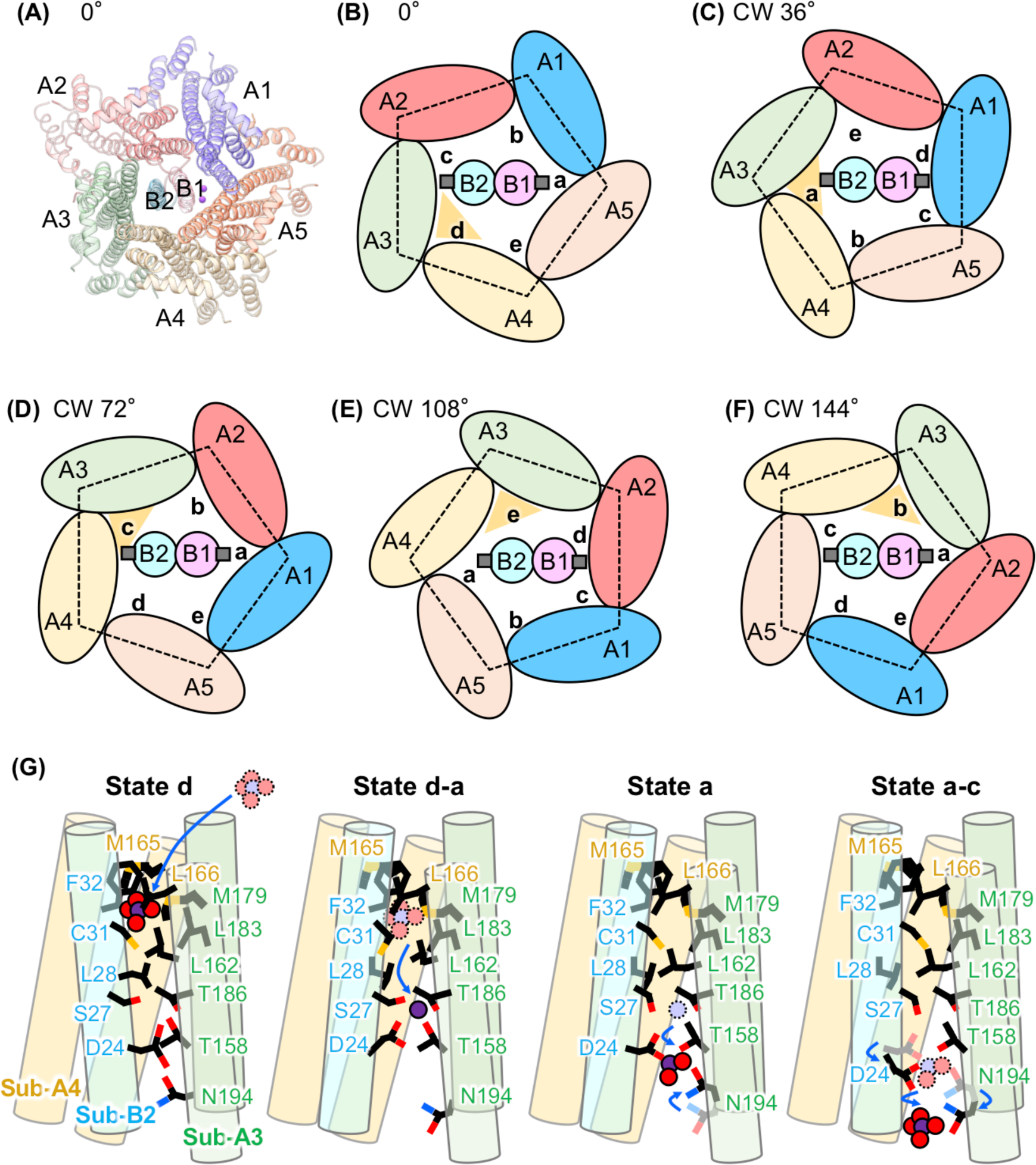
A model of sodium ion flux coupled with PomA rotation. **(A-F)** Schematic diagram of the sodium ion flux coupled with PomA rotation viewed from the periplasm. The subunits are painted in the same colors as in Fig. 1D and labeled with capital letters. The structural states of the clefts between two PomA subunits are labeled with small letters. The clefts transition their structural states d, a, c, e, and b with every 36° CW rotation in this order. A sodium ion is imported into Cavity-I’ of the cleft at state d, moves to Cavity-II at state a, and is released to the cytoplasm at state c. The clefts at state b and e do not contain sodium ions. **(A)** Ribbon representation of *Va*-PomAB. **(B-F)** Schematic diagram focused on the cleft between Sub-A3 and A4 (the A3A4-cleft) highlighted by an orange triangle. **(B)** Schematic representation of *Va*-PomAB shown in **(A)**. This subunit arrangement is defined as 0°. A hydrated sodium ion is imported into Cavity-I’ of the A3A4-cleft. **(C)** After 36° CW rotation of the PomA ring, the state of the A3A4-cleft transitions to state a, and the hydrated sodium ion moves to Cavity-II. The sodium ion is dehydrated by S27 and C31 of PomB and bound to T158 and T186 of PomA. **(D)** After 72° CW rotation of the PomA ring, the state of the A3A4-cleft transitions to state c. The sodium ion is picked by D24 of PomB and is released to the cytoplasm via N194. **(E)** The state of the clefts after 108° CW rotation of the PomA ring. **(F)** The state of the clefts after 144° CW rotation of the PomA ring. **(G)** Side view of the schematic diagram of the sodium ion flux through the A3A4-cleft. For easy viewing, Sub-B2 is rotated relative to the PomA subunits. Sodium ion and hydrated waters are shown by purple and red balls, respectively.

A hydrated sodium ion would be imported into Cavity-I’ at state d or during the transition from b to d because no cavity corresponding to Cavity-I is formed at state b, c, and e, and state a is next to state d. The periplasmic side of Cavity-I’ is closed, even if the plug region is removed from the model. Thus, the entrance for Na^+^ is still unknown. The structures of MotAB from *Campylobacter jejuni* and its plug deletion variant revealed that removal of the plug induces only small changes in the stator structure (*21*). Therefore, the sodium entrance may open during the transition from b to d. The hydrated sodium ion stays in Cavity-I’ at state d because the cytoplasmic side of Cavity-I’ is closed. Along with the CW rotation of PomA, the pathway from Cavity-I’ to Cavity-II opens and D24 is detached from T158 and T186. After 36 degrees of CW rotation, the state of the cleft transitions to state a (**Fig. 6**). Then the hydrated sodium ion moves to Cavity-II and interacts with S27 and C31 of PomB, and T158 and T186 of PomA. The sodium ion may be dehydrated by these residues and bound to T158 and T186 as shown by Hu et al (*22*). A study of the *V. cholerae* stator reported that S26 of *V. cholerae* PomB, which corresponds to S27 of *Va-*PomB, is involved in the dehydration of sodium ions (*40*). PomA and PomB of *V. cholerae* show 89.3% and 86.4 % amino acid sequence identity to those of *V. alginolyticus*. The residues on the tunnel wall are fully conserved between *V. cholerae* and *V. alginolyticus*. Therefore, S27 may contribute to the dehydration of sodium ions in the *Va-*stator. C31 of PomB may also contribute to the dehydration of sodium ion, because C31S and C31T mutations show no effect on motility whereas substitution of C31 to G, A, V, or I reduced the motility and to M conferred non-motile phenotype (*41*). The sodium ion bound to T158 and T186 is then picked by D24 and released to the cytoplasm during the transition from a to c. N194 of PomA may be involved in this process because the PomA N194D mutation suppresses the PomB D24N mutation (*42*). N194 comes close to D24 by slight CW rotation from state a, and the sodium ion bound to D24 may be passed to N194 before release to the cytoplasm.

The PomAB stator conducts sodium ions but not protons. How does the stator discern the specific ions? Our structure suggests that Cavity-I is involved in ion selectivity as a size filter, and the following mutation analysis is consistent with this idea. The amino acid sequence comparison between the sodium and proton-type stators also supports the idea (fig. S3). Among the nine residues forming Cavity-I, seven residues are fully conserved in the Na^+^-type stators. These residues are well conserved among the H^+^-type stators but replaced by residues smaller than those of the Na^+^-type stators. The M165 and M179 of PomA are fully conserved in the Na^+^-type stators but replaced by alanine for M165 and isoleucine for M179 in H^+^-type stators. PomB L35 conserved in the Na^+^-type stators are valine in most H^+^-type stators.

Hu et al proposed that T158, T185, and T186 of PomA are part of the PomAB ion selectivity filter, based on the structure of PomAB at 300 mM NaCl and sequence comparison between sodium and proton-type stators (*22*). However, these residues are also conserved in the H^+^-type MotAB stator of *Rhodobacter sphaeroides* (*Rs*-MotAB) (fig. S3). The *Rs*-MotAB is highly homologous to *Va*-PomAB (*5,43,44*) with a sequence identity of 40 % for the A-subunit and 55 % for the transmembrane region of the B-subunit. Moreover, the residues in the putative ion pathway are conserved except for three B-subunit residues, S27, L28, and C31, which are located in the bottleneck region between Cavity-I and -II (fig. S3). Mutation of L28 and C31 may affect the size and hydrophobicity of Cavity-I, and S27 and C31 may be involved in the dehydration of the ion. These facts suggest that ion selectivity is not determined by a few key residues but by a balance of multiple factors including Cavity-I, the residues involved in the dehydration of ions in the bottleneck region, and the threonine residues in Cavity-II.

Our structure model of *Va*-PomAB in complex with phenamil is consistent with a previous study on chimeric stator between *Va*-PomAB and *Rs*-MotAB (*45*). The stator consisting of PomA and a chimeric B-subunit MomB6 (N-terminal 23 residues of *Va*-PomB are replaced by N-terminal 38 residues of *Rs*-MotB) is driven by sodium and resistant to phenamil. Comparing the amino acid sequence between G14 and F22 of *Va*-PomB with *Rs*-MotA, there are four different residues (L15A, L17A, M19L, G20A) (fig. S3). These residues are involved in phenamil binding in our structure. L15 directly interacts with the pyrazine ring of phenamil. L17 is a part of the short narrow tunnel accommodating the Cl atom of phenamil, and M19 is a part of the hydrophobic pocket for the phenyl group of phenamil. G20 is in contact with phenamil, and if G20 is replaced by alanine, steric hindrance will occur. Therefore, the replacement of these four residues considerably reduced the binding affinity for phenamil.

F22Y mutation of PomB confers phenamil resistance on the stator (*46*). F22 of Sub-B1 is a part of the hydrophobic pocket for the phenyl group of phenamil, and the hydrophobic pocket may be collapsed by substitution with tyrosine. Another previous mutation study revealed that D148 of PomA and P16 of PomB contribute to phenamil binding (*32*). Although D148 of PomA is near phenamil, P16 is a little bit far from phenamil. If the PomA ring slightly rotates in the CW direction, P16 comes close to phenamil. Phenamil may be loosely bound in our structure. The weak density of phenamil in our map is probably due to the loose binding of phenamil.

In this study, we proposed a model of sodium ion flux coupled with the PomA ring rotation around PomB, but the mechanism of how the ion flux drives the rotation is still unknown. PomB D24, the key residue of ion conductivity, did not bind sodium ions in our structure. The side chain conformation of D24 is almost the same as that of the PomB-D24N mutant, which does not conduct sodium ions. Therefore, the actual role of D24 on stator function including ion conductivity is still unclear. D24 would bind sodium ions in a different relative rotation angle of the A-subunit ring. Up to now, 12 stator structures, including our five structures, have been deposited in PDB. However, most of the structures are determined with a ‘plug’, and only one ‘unplugged’ structure is reported. All known structures, even the unplugged structure, have almost the same relative rotation angle of the A-subunit ring, and thus they are basically in the same state. To fully understand the driving mechanism of the stator coupled with ion flow, we need the stator structures with various relative rotational geometry between the A-subunit ring and the B-subunit dimer.

## Materials and Methods

### Bacterial strains and plasmids

The bacterial strains and the plasmids used in this study are listed in table S1. *E. coli* cells were cultured in LB medium (Lennox, Nacalai tesque). If needed, ampicillin or chloramphenicol were added at a final concentration of 50 or 25 μg mL^-1^ each. *V. alginolyticus* cells were cultured in VC medium (0.5% [w/v] polypeptone, 0.5% [w/v] yeast extract, 0.4% [w/v] K2HPO4, 3% [w/v] NaCl, 0.2% [w/v] glucose) or VPG medium (1% [w/v] polypeptone, 0.4% [w/v] K2HPO4, 3% [w/v] NaCl, 0.5% [w/v] glycerol). If needed, chloramphenicol was added at a final concentration of 2.5 μg mL^-1^. Site-directed mutagenesis was performed using the QuikChange method (Agilent Technologies). Transformation of *E. coli* was performed using a standard heat shock method.

### Purification of the stator

Purification and detection of the *Va*-PomAB stator were carried out by the method previously described (*30*). BL21(DE3)/pLysS cells carrying the plasmid pColdIV-*pomApomB*-*his6* with no mutation (wild-type) or with *the pomB*-*D24N* mutation were grown overnight in 20 mL of LB medium at 37°C, inoculated in 2 L of LB medium, and cultured at 37°C to an optical density at 660 nm of 0.5. After incubation on ice for 30 min, IPTG was added to a final concentration of 0.5 mM and cultured for 1 day at 15°C. The cells were pelleted by centrifugation at 8,000 × *g* for 10 min and suspended in 7 mL of Na-Pi buffer (50 mM sodium phosphate [pH 8.0], 200 mM NaCl, and 10% [w/v] Glycerol) per 1 g (wet weight) of the cells. The suspension was passed through a French press (5501-M, Ohtake Works) twice at 1,000 kg cm^-2^. After the removal of unbroken cells by low-speed centrifugation at 8,000 × *g* for 10 min, the cell lysate was ultra-centrifuged at 118,000 × *g* for 1 h. The pellet was suspended in the original volume of Na-Pi buffer and stored at −30°C until use.

The frozen suspension was thawed in a water bath, solubilized in 5% (w/v) lauryl maltose neopentyl glycol (2,2-didecylpropane-1,3-bis-β-D-maltopyranoside, LMNG) in double distilled water to the final concentration of 0.5% (w/v), and stirred for at least 60 min at 4°C. Insoluble materials were removed by centrifugation at 120,000 × *g* for 30 min. The supernatant was subjected to HisTrap FF 5 mL column (Cytiva) equilibrated with buffer A (50 mM Na-Pi [pH 8.0], 200 mM NaCl, 25 mM Imidazole 10% [w/v] Glycerol and 0.01% [w/v] LMNG), and proteins were eluted with a linear gradient of 25−250 mM imidazole produced by mixing buffer A with buffer B (50 mM Na-Pi [pH 8.0], 200 mM NaCl, 500 mM Imidazole, 10 % [w/v] Glycerol and 0.01% [w/v] LMNG). Peak fractions containing the stator complex were concentrated to 500 µL using a 100 K Amicon device (Millipore), loaded on the size-exclusion column (ENrich SEC650 10/300 column, Bio Rad) equilibrated with buffer C (20 mM Tris HCl [pH8.0], 100 mM KCl or NaCl and 0.0025% [w/v] LMNG) and eluted with buffer C at flow rate of 0.75 mL min^-1^. The peak fractions were collected and concentrated using an Amicon device with a 100 kDa cutoff (Millipore). The concentration of the purified sample was measured by 280 nm absorption using nano-drop (Thermo Scientific). LMNG in buffer A, buffer B, and buffer C was replaced by glyco-diosgenin (GDN) (0.5%, 0.01%, and 0.005% [w/v] GDN for buffer A, buffer B, and buffer C, respectively) for purification of *Va*-PomAB used for the preparation of the phenamil complex. The purity of the sample was assessed by sodium dodecyl sulfate polyacrylamide gel electrophoresis (SDS-PAGE) with Coomassie Brilliant Blue R250 (CBB) staining.

### Cryo-EM data collection

10 µL of purified *Va*-PomAB-His6 solution (1.49 mg mL^-1^ for 100 mM NaCl sample and 3.46 mg mL^-1^ for 100 mM KCl sample) or *Va*-PomAB(D24N)-His6 solution (3.25 mg mL^-1^ for 100 mM NaCl sample and 3.04 mg mL^-1^ for 100 mM KCl sample) were applied to a Quantifoil holey carbon grid R1.2/1.3 or R0.6/1.0 Cu 300 mesh (Quantifoil Micro Tools GmbH, Großlöbichau, Germany) with pretreatment of a side of the grid by glow discharge. 3 µL of freshly purified protein was applied to the grids, and the grids were plunged into liquid ethane for rapid freezing with Vitrobot Mark IV (Thermo Fisher Scientific) with a blotting time of 4 - 7 s at 4 °C and 100% humidity.

The cryoEM sample of *Va*-PomAB-His6 with phenamil was prepared by mixing 99 µL of *Va*-PomAB-His6 (6.67 mg mL^-1^) purified using GDN with 1 µL of DMSO solution of 10 mM phenamil methanesulfonate (Sigma) to a final concentration of 100 µM Phenamil, 1% (w/v) DMSO, and 6.60 mg mL^-1^ *Va*-PomAB-His6, followed by about 10 min incubation on ice. 3 µL of the sample solution was applied to a Quantifoil holey carbon grid R1.2/1.3 and R0.6/1.0 Cu 300 mesh with pretreatment of a side of the grid by glow discharge. The grids were plunged into liquid ethane for rapid freezing with Vitrobot Mark IV (Thermo Fisher Scientific) with a blotting time of 5 - 6 s at 4 °C and 100% humidity.

The cryoEM data were collected using a Titan Krios electron microscope (FEI) equipped with a thermal field-emission electron gun operated at 300 kV, a Gatan energy filter, and a K3 direct electron detector camera (Gatan, USA). The cryoEM movie data were automatically collected using the SerialEM software. The dose-fractionated movies were taken at a nominal magnification of 105,000×, corresponding to an image pixel size of 0.675 Å. We collected four image data sets (1,248 (set-1), 3,128 (set-2), 5,542 (set-3), and 7,181 (set-4) micrographs) for *Va-*PomAB with NaCl, two image data sets (3,947 (set-1) and 13,730 (set-2) micrographs) for *Va-*PomAB with KCl, two image data sets (5,450 (set-1) and 2,717 (set-2) micrographs) for *Va-*PomAB(D24N) with NaCl, four image data sets (1,822 (set-1), 3,531 (set-2), 3,891 (set-3) and 4,307 (set-4) micrographs) for *Va-* PomAB(D24N) with KCl, and three image data sets (2,548 (set-1), 5,808 (set-2), and 5,518 (set-3) micrographs) for *Va-*PomAB with phenamil and NaCl. CryoEM data collection is summarized in table S2.

### Image Processing

All micrographs were motion-corrected using MOTIONCORR2 (version 1.3.0) with dose fraction (*47*), and CTF values were estimated with CTFFIND 4.1 (version 1.10) (*48*). The image processing was performed using RELION-3.1 (*49*) and 4β (*50*). The image processing schemes are shown in fig. S4 to S8.

The image processing of *Va-*PomAB with NaCl was initiated by manual picking of 1805 particles from 54 micrographs in set-1. The particle images were 2D class averaged, and 507 good particle images were selected as templates for auto-picking with the template matching procedure. 345,375 particles auto-picked from 1,248 micrographs of set-1 were 2D class averaged, and 19,655 particles were selected as second templates for auto-picking. 274,999 auto-picked particles were 2D and 3D class averaged, and 30,697 good particle images were used for the training data set for Topaz auto-picking. 127,822 particle images were picked from 1,248 micrographs of set-1 by Topaz auto-picking and were 2D-classified. 63,671 good particle images were applied for 3D classification using the density map of *Aquifex aeolicus* MotA pentamer (*23*), and 44,067 good particle images were used for 3D refinement, polish, and CTF refinement. On the other hand, 465,830, 1,685,742, and 693,855 particles were extracted from 3,138 micrographs of set-2, 5,542 micrographs of set-3, and 7181 micrographs of set-4 by Topaz auto-picking, respectively, and false particles were removed by two rounds of 3D classification. After 3D refinement, polish, and CTF refinement, 140,531 remaining particles from set-2 were combined with the 44,067 particles from set-1 and applied for 3D classification to select 177,048 good particles. After 3D refinement, polish, and CTF refinement, 303,302 remaining particles from set-3 were combined with the 177,048 particles from set-1 and -2 and applied for 3D classification to select 382,159 good particles. After 3D refinement, polish, and CTF refinement, 279,856 remaining particles from set-4 were combined with the 382,159 particles from set-1, -2, and -3 and applied for 3D refinement followed by postprocess. The final cryo-EM map showed a 3.1 Å resolution (fig. S9).

The image processing of *Va-*PomAB with KCl was started by auto-picking of 1,237,150 particles from 3,947 micrographs of set-1 with the template matching procedure using the density map of *Va-*PomAB with NaCl. After 2D class averaging, 52,600 good particle images were used for the training data set for Topaz auto-picking. 2,220,744 particle images were auto-picked from 3,947 micrographs of set-1 by Topaz and 2D-averaged. 55,426 good particle images were applied for 3D classification using the density map of *Va-*PomAB with NaCl. 37,175 good particle images were used for 3D refinement, polish, and CTF refinement, and the density map at 4.2 Å resolution was obtained. Meanwhile, 13,730 micrographs of set-2 were separated into two subsets, subset-1 (3,433 micrographs) and subset-2 (10,297 micrographs), and 1,896,621 particles were auto-picked from subset-1with template matching procedure using the density map of *Va-* PomAB with NaCl. After 2D class averaging. 636,595 good particle images were 2D and 3D class averaged, and 65,475 good particle images were used for the training data set for Topaz auto-picking. Using Topaz auto-picking, 599,146 and 1,801,795 particles were extracted from subset-1 and -2, respectively, and false particles were removed by three rounds of 3D class averaging. Then the good particles from subset-1 and -2 were merged, and 292,671 particles were 3D class-averaged after 3D refinement. 246,999 good particle images were used for 3D refinement, polish, and CTF refinement, and the density map at 3.7 Å resolution was obtained. Then, the 246,999 particle images were merged with the 37,175 particle images from set-1 used for the 4.2 Å resolution map construction. After 3D class averaging, 255,000 good particle images were applied for 3D refinement, polish, and CTF refinement. We finally obtained a density map at 3.4 Å resolution after 3D refinement and postprocess (fig. S9).

The micrograph data set-1 of *Va-*PomAB(D24N) with NaCl was separated into three subsets, subset-1 (765 micrographs), subset-2 (4,327 micrographs), and subset-3 (358 micrographs), and 369,110 particles were auto-picked from subset-1with template matching procedure using the density map of *Va-*PomAB with NaCl. After 2D and 3D class averaging, 19,104 good particle images were used for the training data set for Topaz auto-picking. Using the Topaz auto-picking, 49,990, 374,009, and 45,333 particle images were auto-picked from micrographs of subset-1, -2, and -3, respectively, and the particle images from subset-1 and -2 were merged. After 3D class averaging using the density map of *Va-*PomAB with NaCl, 227,695 good particle images from subset-1 and -2 were merged with the 25,359 good particle images from subset-3 which were obtained after two rounds of 3D class averaging, and the merged 253,054 good particle images were applied for 3D refinement, polish, and CTF refinement. Meanwhile, 151,848 particle images were auto-picked from 2,717 micrographs of set-2 by Topaz. After two rounds of 3D class averaging, 735,221 good particle images were applied for 3D refinement, polish, and CTF refinement. Then the 253,054 particle images from set-1 and 735,221 particle images were merged and applied for 3D refinement, CTF refinement, and postprocess. The final cryo-EM map showed a 3.2 Å resolution (fig. S9).

The image processing of *Va-*PomAB(D24N) with KCl was begun by manual picking of 2,218 particles from 80 micrographs in set-1. After 2D class averaging, 617 good particle images were selected for auto-picking with the template matching procedure. 412,521 particles auto-picked from 1,822 micrographs of set-1 were 2D class averaged, and 65,987 good particles were used for the training data set for Topaz auto-picking. Using the Topaz auto-picking, 92,440, 158,767, 219,813, and 253,069 particle images were auto-picked from micrographs of set-1, -2, -3, and -4, respectively. After two rounds of 3D class averaging, 46,088, 86,282, 137,208, and 143,562 good particles were selected from set-1, -2, -3, and -4, respectively, applied for 3D refinement and polish, and merged. 3D class averaging was performed for the merged data, and 325,940 good particles were applied for 3D refinement, CTF refinement, and postprocess. The final cryo-EM map showed a 3.4 Å resolution (fig. S9).

The image processing of *Va-*PomAB with phenamil was started by manual picking of 3,902 particles from 39 micrographs in set-1. The particle images were 2D class averaged, and 1,061 good particle images were selected as templates for auto-picking with the template matching procedure. 567,034 particles auto-picked from 2,548 micrographs of set-1 were 2D class averaged followed by two rounds of 3D class averaging, and 68,937 good particle images were used for the training data set for Topaz auto-picking. 605,249 particle images were picked from set-1 by Topaz and were 3D-classified. 111,573 good particle images were further 3D classified, and 44,816 good particle images were used for 3D refinement, polish, and CTF refinement. The data set-2 was separated into two subsets, subset-1 (3,260 micrographs) and subset-2 (2,548 micrographs). 1,282,092 particles were auto-picked from subset-1, and 137,275 good particles selected after three rounds of 3D classification were applied for 3D refinement, polish, and CTF refinement. 1,128,968 particles were auto-picked from subset-2, and 221,533 good particles selected after three rounds of 3D classification were applied for 3D refinement, polish, and CTF refinement. The final good particles of subset-1 and -2 from set-2 were merged with the final good particles from set-1. After 3D classification, 346,168 good particles from the merged data were applied for 3D refinement and CTF refinement. 1,556,863 particles were auto-picked from set-3. 250,734 good particles selected after four rounds of 3D classification were applied for 3D refinement, polish, and CTF refinement, and combined with the 346,168 good particle data from set-1 and -2. Then the combined 596,902 particles were applied for 3D refinement, CTF refinement, and postprocess. The final cryo-EM map showed a 3.3 Å resolution (fig. S9).

### Model building and refinement

The atomic models were constructed by Coot (*51*) and refined using Phenix (*52*). Before model building, the cryo-EM map was auto-sharpened by Phenix (*52*). The refinement statistics are summarized in table S2.

### Structural analysis

The structural comparison and analysis were performed using Pymol (Schrödinger LLC), Chimera (*53*), and Chimera X (*54*). The analysis of cavities in the stator was performed using CAVER Analyst 2.0 BETA (*55*).

### Measurement of cell growth

Overnight cultures of DH5α cells carrying the plasmid pBAD33 or pTSK37 with pomA-L166K, M179K, L166A, or M179A mutation or without mutation at 37°C in LB medium were diluted 100-fold into LB medium or LB-Na0 medium (1% [w/v] bactotryptone, 0.5% [w/v)] yeast extract), and cells were grown at 37°C for 8 h. Arabinose was added from the beginning of the second culture at a final concentration of 0.2% (w/v) for over-production of plug deletion stators. Cell growth was monitored at the optical density of 600 nm (OD600) every hour.

### Amino acid sequence alignment

Amino acid sequences of MotA^Cj^, MotA^Bs^, MotA^Ec^, MotA^Se^, MotA^Rs^, PomA^SO^, PomA^Vp^, PomA^Vm^ and MotP^Bs^ (Cj, *C*. *jejuni*; Bs, *B*. *subtilis*; Ec, *E*. *coli*; Se, *S*. *enterica*; Rs, *R*. *sphaeroides* (as *C*. *sphaeroides*); So, *Shewanella oneidensis*; Vp, *V*. *parahaemolyticus*; Vm, *V*. *mimicus*.) were downloaded from NCBI web sites (https://www.ncbi.nlm.nih.gov/) as the FASTA formats (ID: QMU89816.1, SUY61260, 1QGU24922.1, NP_416404.1, AAC45265.1, EGJ21703.1, WP_006085155.1, WPD15183.1, EEW10820.1 and NP_390851.1, respectively). Amino acid sequences of MotB^Cj^, MotB^Bs^, MotB^Ec^, MotB^Se^, MotB^Rs^, PomB^So^, PomB^Vp^, PomB^Vm^ and MotS^Bs^ were downloaded from NCBI web sites (https://www.ncbi.nlm.nih.gov/) as the FASTA formats (ID: QMU89815.1, SUY61262.1, QGU24923.1, NP_416403.1, AAC45266.1, EGJ21702.1, AAN54590.1, WPD15184.1, EEW10819.1 and NP_390850.1, respectively). The sequence alignment was performed using the ClustalOmega website (https://www.ebi.ac.uk/Tools/msa/clustalo/).

## Data and materials availability

All the data obtained in this study are available in the main text, the Supplementary Data, and the Supplementary Table. The atomic coordinate has been deposited in Protein Data Bank, www.pdb.org under accession code 8ZYV (*Va-* PomAB with 100 mM NaCl), 8ZYW (*Va-*PomAB with 100 mM KCl), 8YZY (*Va-* PomAB (D24N) with 100 mM NaCl), 8ZZ0 (*Va-*PomAB (D24N) with 100 mM KCl), and 9IJM (*Va-*PomAB with phenamil). The cryo-EM maps have been deposited in the Electron Microscopy Data Bank under accession code EMD-60580 (*Va-*PomAB with 100 mM NaCl), EMD-60581 (*Va-*PomAB with 100 mM KCl), EMD-60584 (*Va-*PomAB (D24N) with 100 mM NaCl), EMD-60585 (*Va-*PomAB (D24N) with 100 mM KCl), and EMD-60636 *Va-*PomAB with phenamil).

## Acknowledgments

We thank Akio Kitao, Yohei Miyanoiri and Tran P. Duy for valuable discussions.

## Author contributions

Conceptualization: TN, M. Homma, KI

Methodology: TN, JK, M.Hirose, SK, TK, KI

Investigation: TN, NT, JK, SK, KI

Visualization: TN, NT, KI

Supervision: TN, M.Homma, KI

Writing—original draft: TN, NT, KI

Writing—review & editing: All the authors

