## Supplemental Fig.S1-S9, Table S1 and S2 for "Structural insight into sodium ion pathway in the bacterial flagellar stator from marine *Vibrio*"

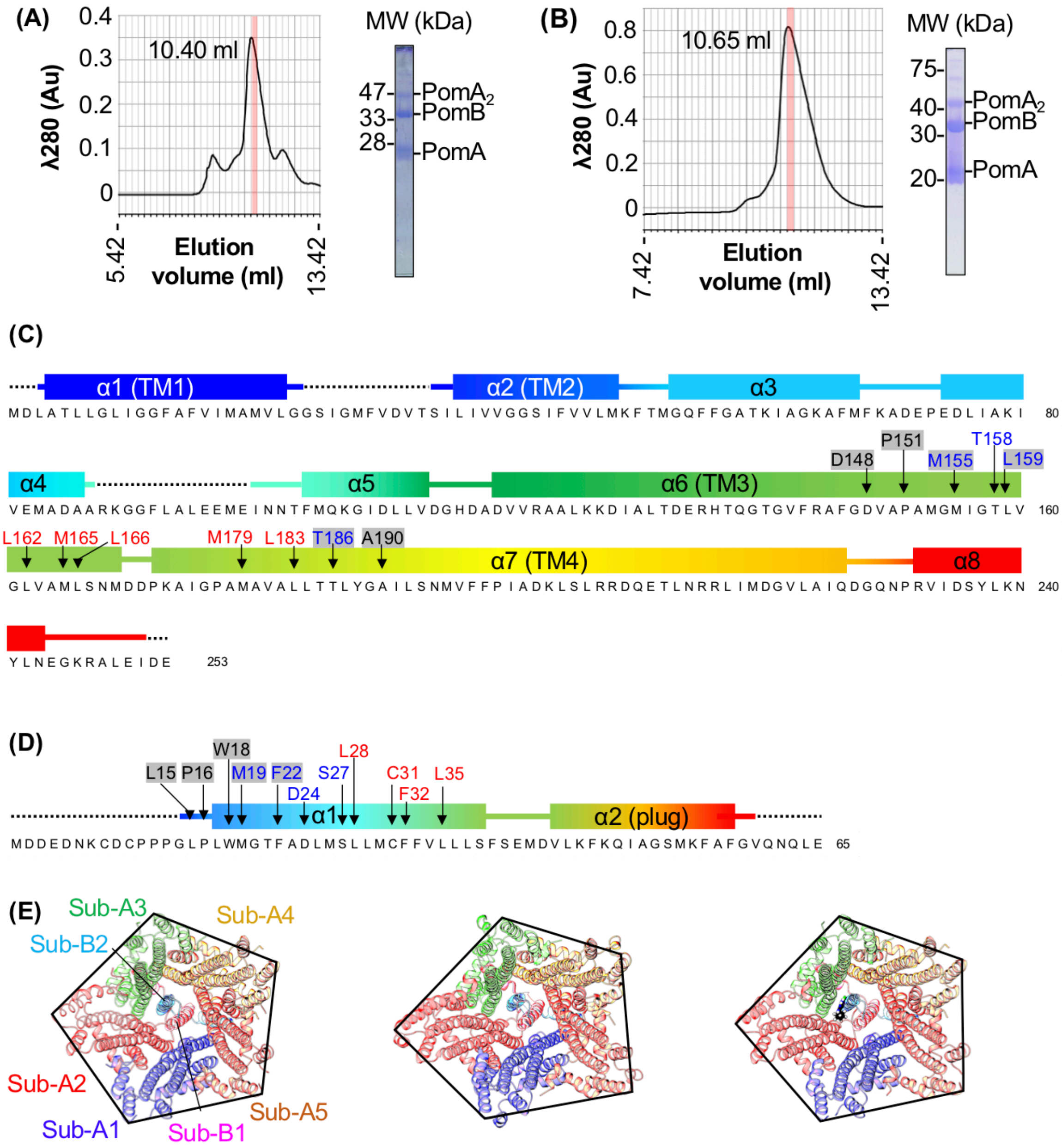


**Fig. S1. Structure analysis of *Va*-PomAB.**

(A) (B) SEC elution profile and SDS-PAGE pattern of *Va*-PomAB (A) and *Va*-PomAB(D24N) (B) samples with LMNG and 100 mM NaCl. The fractions shown by red boxes in the SEC profiles were examined by SDS-PAGE and used for cryo-EM structure analysis. (C) (D) Secondly structure and amino acid sequence of *Va*-PomA (C) and the N-terminal region of *Va*-PomB (D). Residues not included in the model are shown as black dot lines. Residues that form Cavity I and II are indicated by red and blue letters, respectively. Phenamil binding sites are shown by letters with a gray shade. (E) Comparison of the subunit arrangement of *Va*-PomAB with 100 mM NaCl (left), *Va*-PomAB with 300 mM NaCl (middle) (PDB code: 8BRD), and *Va*-PomAB with phenamil (right). Ribbon models viewed from the cytoplasm are drawn. The subunits are colored as in Fig. 1(D). The Cα atoms of A86 (the C-terminus of α4) in the neighboring subunits are linked.

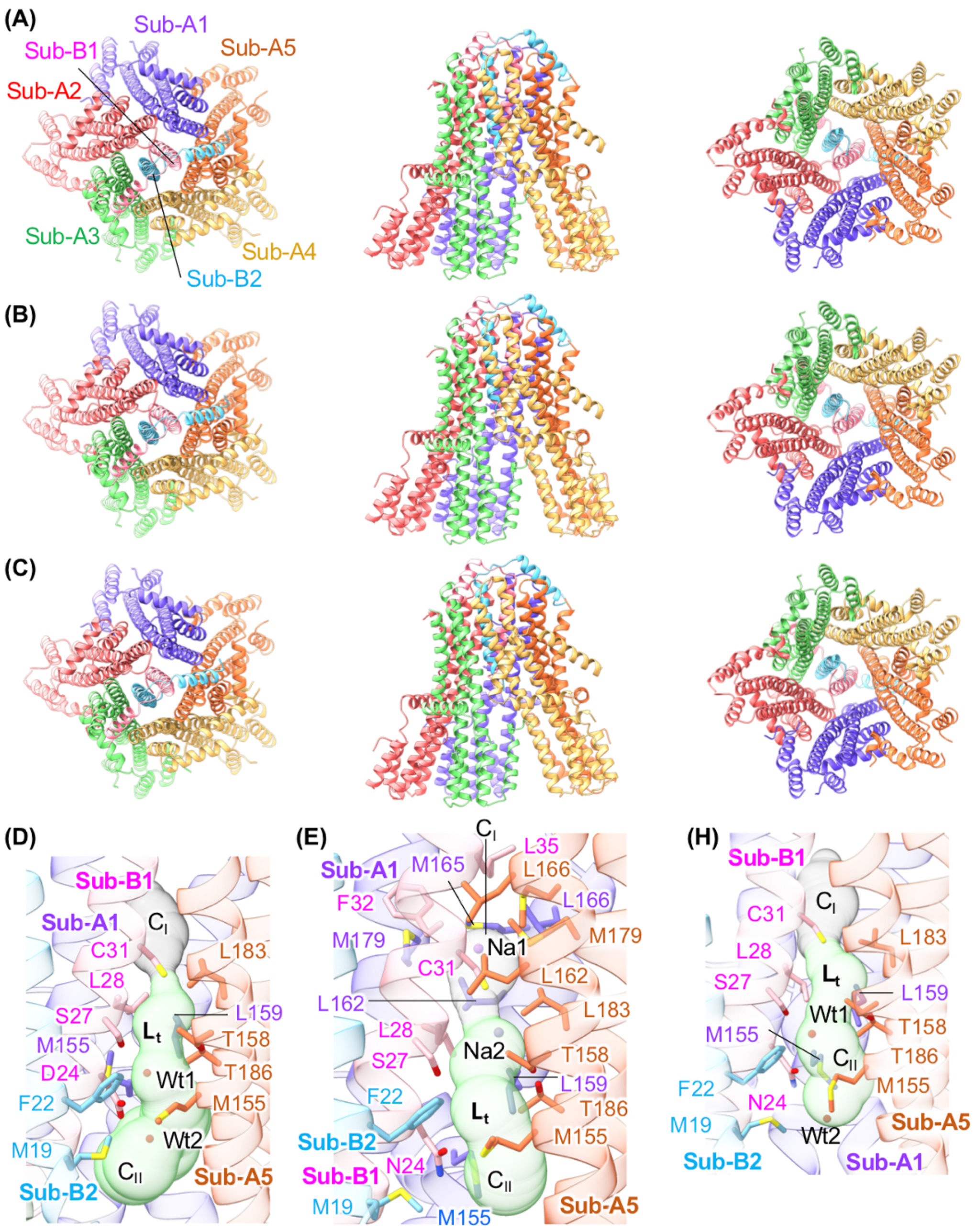


**Fig. S2. Structures of *Va*-PomAB at 100 mM KCl and *Va*-PomAB (D24N) at 100 mM NaCl and at 100 mM KCl.**

Ribbon representation of (A) *Va*-PomAB at 100 mM KCl, (B) *Va*-PomAB (D24N) at 100 mM NaCl, and (C) *Va*-PomAB (D24N) at 100 mM KCl viewed from periplasm (left), side (middle), and cytoplasm (right). Close-up side view of the long tunnel ($L_t$) of (D) *Va*-PomAB at 100 mM KCl, (E) *Va*-PomAB (D24N) at 100 mM NaCl, and (H) *Va*-PomAB (D24N) at 100 mM KCl. The residues that form the inner wall of the long tunnel are shown by the stick model. Sodium ions (Na1 and Na2) and solvent molecules (Wt1 and Wt2) in the long tunnel are indicated by balls. Cavity-I ($C_I$) and Cavity-II ($C_{II}$) are indicated by gray and green blobs, respectively. The subunits are colored as in Fig. 1(D).

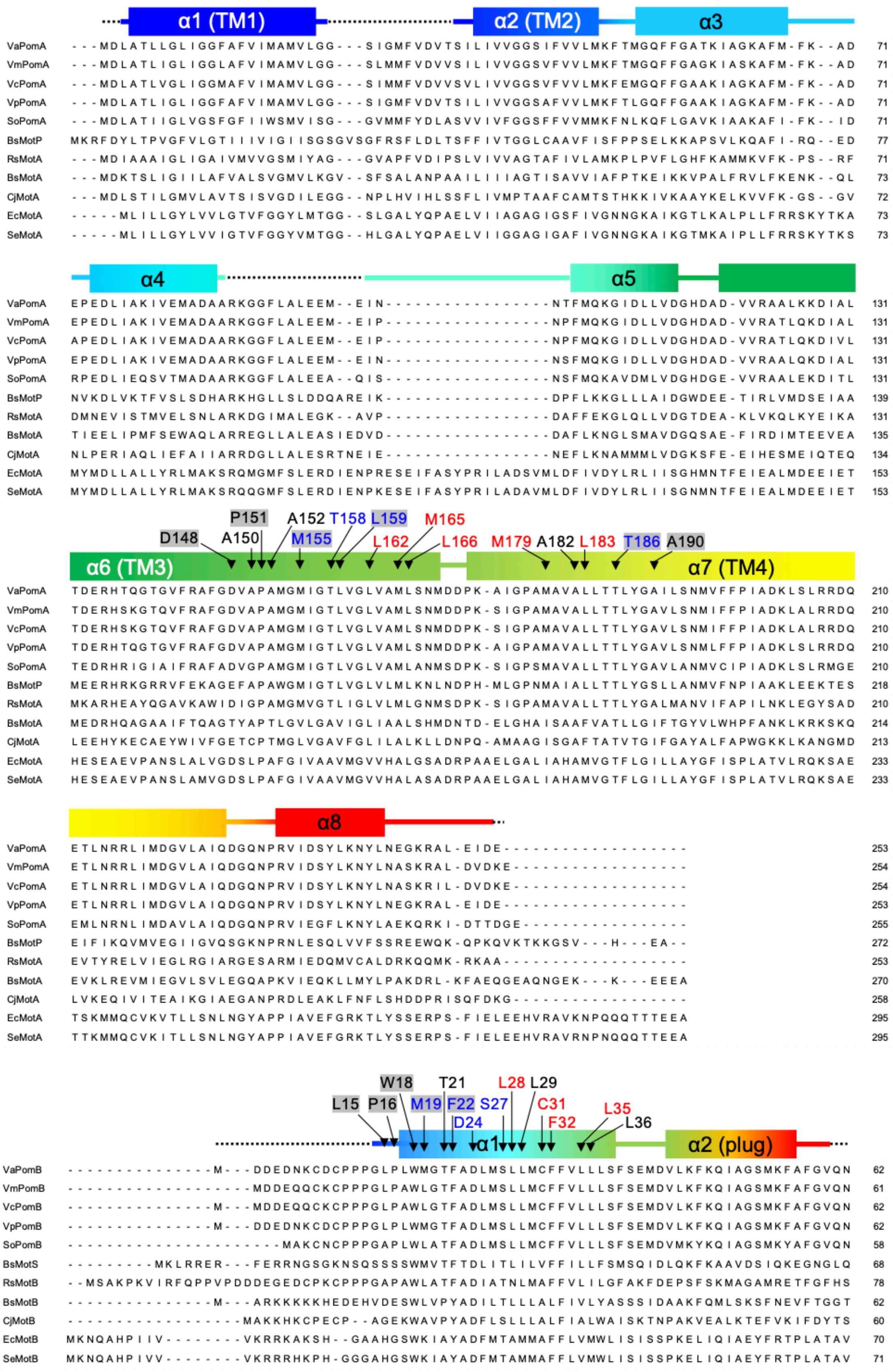

α1 (TM1)
α2 (TM2)
α3
VaPomA ---MDLATLLGLIGGFAFVIMAMVLGG---SIGMFVDVTSILIVVGGSIFVVLMKFTMGQFFGATKIAGKAFM-FK--AD 71
VmPomA ---MDLATLIGLIGGLAFVIMAMVLGG---SLMMFVDVVSILIVVGGSVFVVLMKFTMGQFFGAGKIASKAFM-FK--AD 71
VcPomA ---MDLATLVGLIGGMAFVIMAMVLGG---SIMMFVDVVSVLIVVGGSVFVVLMKFEMGQFFGAAKIAGKAFM-FK--AD 71
VpPomA ---MDLATLIGLIGGFAFVIMAMVLGG---SIGMFVDVTSILIVVGGSAFVVLMKFTLGQFFGAAKIAGKAFM-FK--AD 71
SoPomA ---MDLATIIGLVGSFGFIIWSMVISG---GVMMFYDLASVVIVFGGSFFVVMMKFNLKQFLGAVKIAAKAFI-FK--ID 71
BsMotP MKRFDYLTPVGFVLGTIIIVIGIISGSGVSGFRSFLDLTSFFIVTGGLCAAVFISFPPSELKKAPSVLKQAFI-RQ--ED 77
RsMotA ---MDIAAAIGLIGAIVMVVGSMIYAG---GVAPFVDIPSLVIVVAGTAFIVLAMKPLPVFLGHFKAMMKVFK-PS--RF 71
BsMotA ---MDKTSLIGIILAFVALSVGMVLKGV--SFSALANPAAILIIIAGTISAVVIAFPTKEIKKVPALFRVLFKENK--QL 73
CjMotA ---MDLSTILGMVLAVTSISVGDILEGG--NPLHVIHLSSFLIVMPTAAFCAMTSTHKKIVKAAYKELKVVFK-GS--GV 72
EcMotA -----MLILLGYLVVLGTVFGGYLMTGG--SLGALYQPAELVIIAGAGIGSFIVGNNGKAIKGTLKALPLLFRRSKYTKA 73
SeMotA -----MLILLGYLVVIGTVFGGYVMTGG--HLGALYQPAELVIIGGAGIGAFIVGNNGKAIKGTMKAIPLLFRRSKYTKS 73
α4
α5
VaPomA EPEDLIAKIVEMADAARKGGFLALEEM--EIN-----------------NTFMQKGIDLLVDGHDAD-VVRAALKKDIAL 131
VmPomA EPEDLIAKIVEMADAARKGGFLALEEM--EIP-----------------NPFMQKGIDLLVDGHDAD-VVRATLQKDIAL 131
VcPomA APEDLIAKIVEMADAARKGGFLALEEM--EIP-----------------NPFMQKGIDLLVDGHDAD-VVRATLQKDIVL 131
VpPomA EPEDLIAKIVEMADAARKGGFLALEEM--EIN-----------------NSFMQKGIDLLVDGHDAD-VVRAALQKDIAL 131
SoPomA RPEDLIEQSVTMADAARKGGFLALEEA--QIS-----------------NSFMQKAVDMLVDGHDGE-VVRAALEKDITL 131
BsMotP NVKDLVKTFVSLSDHARKHGLLSLDDQAREIK-----------------DPFLKKGLLLAIDGWDEE-TIRLVMDSEIAA 139
RsMotA DMNEVISTMVELSNLARKDGIMALEGK--AVP-----------------DAFFEKGLQLLVDGTDEA-KLVKQLKYEIKA 131
BsMotA TIEELIPMFSEWAQLARREGLLALEASIEDVD-----------------DAFLKNGLSMAVDGQSAE-FIRDIMTEEVEA 135
CjMotA NLPERIAQLIEFAIIARRDGLLALESRTNEIE-----------------NEFLKNAMMMLVDGKSFE-EIHESMEIQTEQ 134
EcMotA MYMDLLALLYRLMAKSRQMGMFSLERDIENPRESEIFASYPRILADSVMLDFIVDYLRLIISGHMNTFEIEALMDEEIET 153
SeMotA MYMDLLALLYRLMAKSRQQGMFSLERDIENPKESEIFASYPRILADAVMLDFIVDYLRLIISGNMNTFEIEALMDEEIET 153
P151 A152 T158 L159 M165
D148 A150 M155 L162 L166 M179 A182 L183 T186 A190
α6 (TM3)
α7 (TM4)
VaPomA TDERHTQGTGVFRAFGDVAPAMGMIGTLVGLVAMLSNMDDPK-AIGPAMAVALLTTLYGAILSNMVFFPIADKLSLRRDQ 210
VmPomA TDERHSKGTQVFRAFGDVAPAMGMIGTLVGLVAMLSNMDDPK-SIGPAMAVALLTTLYGAVLSNMIFFPIADKLALRRDQ 210
VcPomA TDERHSKGTQVFRAFGDVAPAMGMIGTLVGLVAMLSNMDDPK-SIGPAMAVALLTTLYGAVLSNMIFFPIADKLALRRDQ 210
VpPomA TDERHTQGTGVFRAFGDVAPAMGMIGTLVGLVAMLSNMDDPK-AIGPAMAVALLTTLYGAVLSNMLFFPIADKLSLRRDQ 210
SoPomA TEDRHRIGIAIFRAFADVGPAMGMIGTLVGLVAMLANMSDPK-SIGPSMAVALLTTLYGAVLANMVCIPIADKLSLRMGE 210
BsMotP MEERHRKGRRVFEKAGEFAPAWGMIGTLVGLVLMLKNLNDPH-MLGPNMAIALLTTLYGSLLANMVFNPIAAKLEEKTES 218
RsMotA MKARHEAYQGAVKAWIDIGPAMGMVGTLIGLVLMLGNMSDPK-SIGPAMAVALLTTLYGALMANVIFAPILNKLEGYSAD 210
BsMotA MEDRHQAGAAIFTQAGTYAPTLGVLGAVIGLIAALSHMDNTD-ELGHAISAAFVATLLGIFTGYVLWHPFANKLKRKSKQ 214
CjMotA LEEHYKECAEYWIVFGETCPTMGLVGAVFGLILALKLLDNPQ-AMAAGISGAFTATVTGIFGAYALFAPWGKKLKANGMD 213
EcMotA HESEAEVPANSLALVGDSLPAFGIVAAVMGVVHALGSADRPAAELGALIAHAMVGTFLGILLAYGFISPLATVLRQKSAE 233
SeMotA HESEAEVPANSLAMVGDSLPAFGIVAAVMGVVHALASADRPAAELGALIAHAMVGTFLGILLAYGFISPLATVLRQKSAE 233
α8
VaPomA ETLNRRLIMDGVLAIQDGQNPRVIDSYLKNYLNEGKRAL-EIDE------------------- 253
VmPomA ETLNRRLIMDGVLAIQDGQNPRVIDSYLKNYLNASKRAL-DVDKE------------------ 254
VcPomA ETLNRRLIMDGVLAIQDGQNPRVIDSYLKNYLNASKRIL-DVDKE------------------ 254
VpPomA ETLNRRLIMDGVLAIQDGQNPRVIDSYLKNYLNEGKRAL-EIDE------------------- 253
SoPomA EMLNRNLIMDAVLAIQDGQNPRVIEGFLKNYLAEKQRKI-DTTDGE----------------- 255
BsMotP EIFIKQVMVEGIIGVQSGKNPRNLESQLVVFSSREEWQK-QPKQVKTKKGSV---H---EA-- 272
RsMotA EVTYRELVIEGLRGIARGESARMIEDQMVCALDRKQQMK-RKAA------------------- 253
BsMotA EVKLREVMIEGVLSVLEGQAPKVIEQKLLMYLPAKDRL-KFAEQGEAQNGEK---K---EEEA 270
CjMotA LVKEQIVITEAIKGIAEGANPRDLEAKLFNFLSHDDPRISQFDKG------------------ 258
EcMotA TSKMMQCVKVTLLSNLNGYAPPIAVEFGRKTLYSSERPS-FIELEEHVRAVKNPQQQTTTEEA 295
SeMotA TTKMMQCVKITLLSNLNGYAPPIAVEFGRKTLYSSERPS-FIELEEHVRAVRNPNQQQTTEEA 295
W18 T21 L28 L29
L15 P16 M19 F22 S27 C31 L35
D24 F32 L36
α1
α2 (plug)
VaPomB ---------------M---DDEDNKCDCPPPGLPLWMGTFADLMSLLMCFFVLLLSFSEMDVLKFKQIAGSMKFAFGVQN 62
VmPomB -------------------MDDEQQCKCPPPGLPAWLGTFADLMSLLMCFFVLLLSFSEMDVLKFKQIAGSMKFAFGVQN 61
VcPomB ---------------M---MDDEQQCKCPPPGLPAWLGTFADLMSLLMCFFVLLLSFSEMDVLKFKQIAGSMKFAFGVQN 62
VpPomB ---------------M---DDEDNKCDCPPPGLPLWMGTFADLMSLLMCFFVLLLSFSEMDVLKFKQIAGSMKFAFGVQN 62
SoPomB ----------------------MAKCNCPPPGAPLWLATFADLMSLLMCFFVLLLSFSEMDVMKYKQIAGSMKYAFGVQN 58
BsMotS ---------MKLRRER---FERRNGSGKNSQSSSSWMVTFTDLITLILVFFILLFSMSQIDLQKFKAAVDSIQKEGNGLQ 68
RsMotB --MSAKPKVIRFQPPVPDDDEGEDCPKCPPPGAPAWLATFADIATNLMAFFVLILGFAKFDEPSFSKMAGAMRETFGFHS 78
BsMotB ---------------M---ARKKKKKHEDEHVDESWLVPYADILTLLLALFIVLYASSSIDAAKFQMLSKSFNEVFTGGT 62
CjMotB ------------------MAKKHKCPECP--AGEKWAVPYADFLSLLLALFIALWAISKTNPAKVEALKTEFVKIFDYTS 60
EcMotB MKNQAHPIIV--------VKRRKAKSH--GAAHGSWKIAYADFMTAMMAFFLVMWLISISSPKELIQIAEYFRTPLATAV 70
SeMotB MKNQAHPIVV--------VKRRRHKPH-GGGAHGSWKIAYADFMTAMMAFFLVMWLISISSPKELIQIAEYFRTPLATAV 71

**Fig. S3. Amino acid sequence alignment of *Va*PomA and *Va*PomB homologs from different bacteria.**

The secondary structure elements of VaPomA and B are shown above the sequence. Residues not included in the model are shown as black dot lines. Residues that form Cavity-I and II are indicated by red and blue letters, respectively. Phenamil binding sites are shown by letters with a gray shade. The aligned sequences are *Vibrio alginolyticus*, VaPomA and B; *Vibrio mimicus,* VmPomA and B; *V. parahaemolyticus,* VpPomA and B; *Shewanella oneidensis*, SoPomA and B; *Bacillus subtilis*, BsMotP, S, A, and B; *Rhodobacter sphaeroides*, RsPomA and B; *Campylobacter jejuni*, CjMotA and B; *Escherichia coli*, EcMotA and B; *Salmonella enterica*, SeMotA and B.

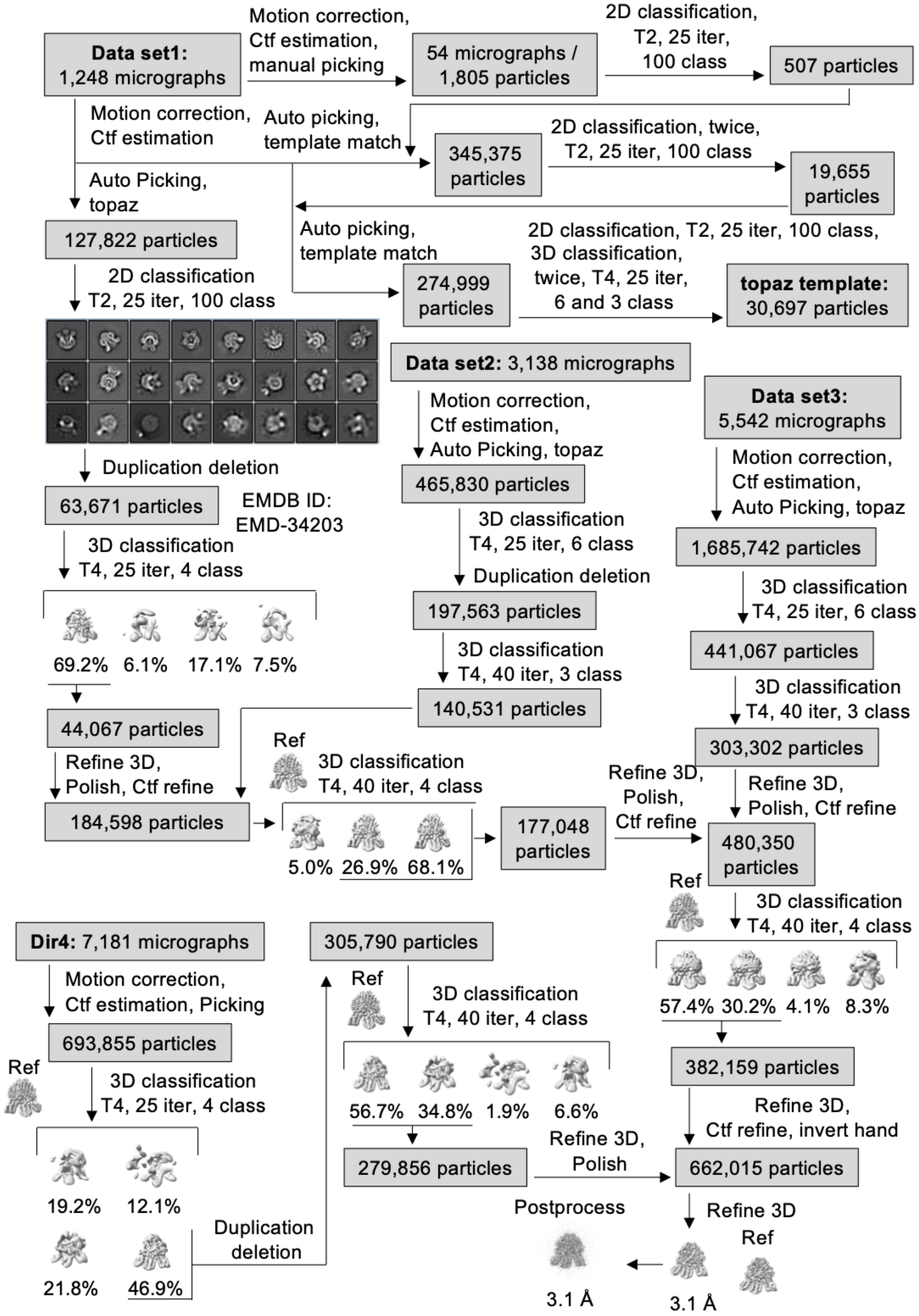


**Fig. S4. 3D reconstruction scheme of *Va*-PomAB with NaCl.**

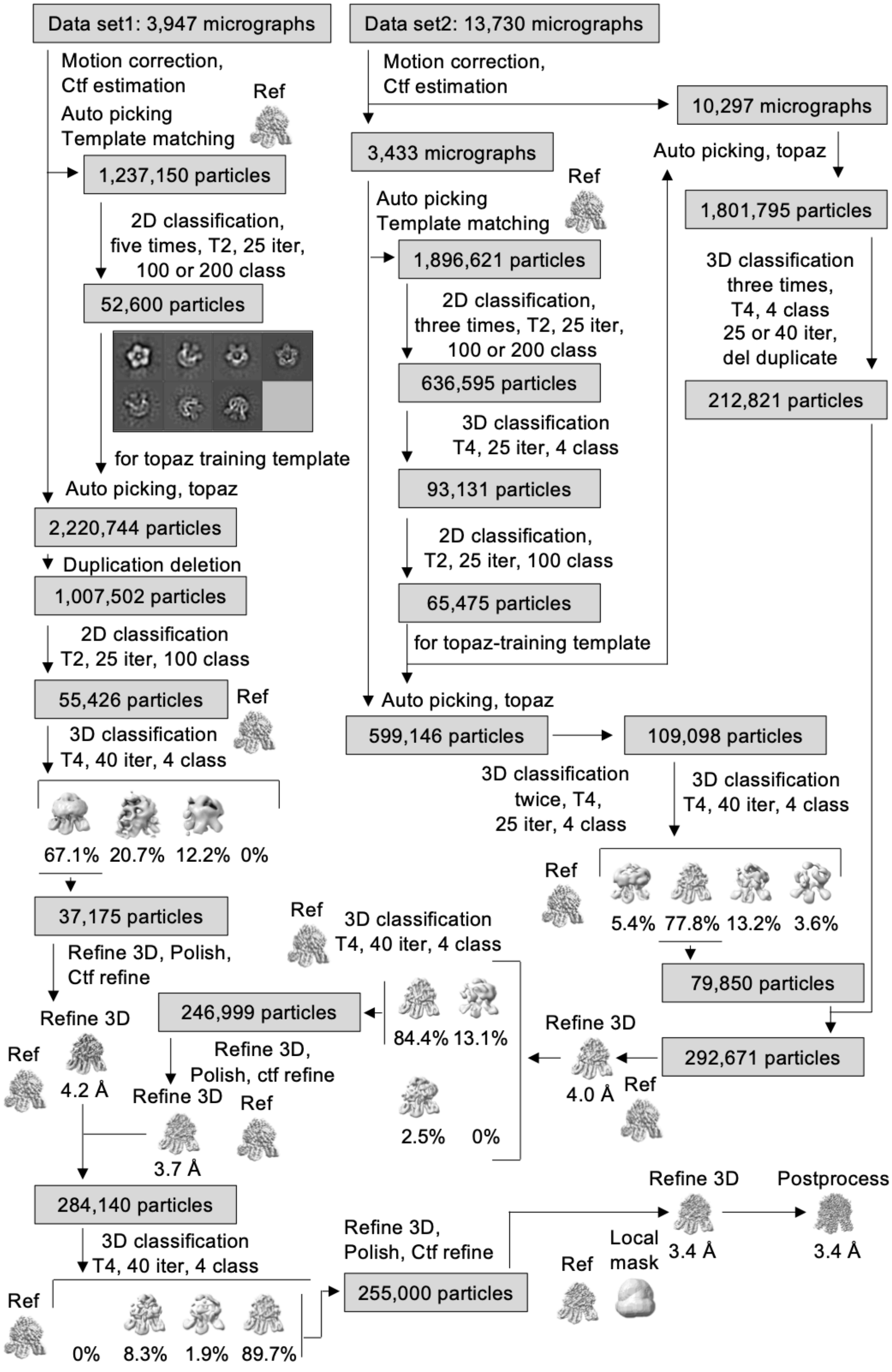


**Fig. S5. 3D reconstruction scheme of *Va*-PomAB with KCl.**

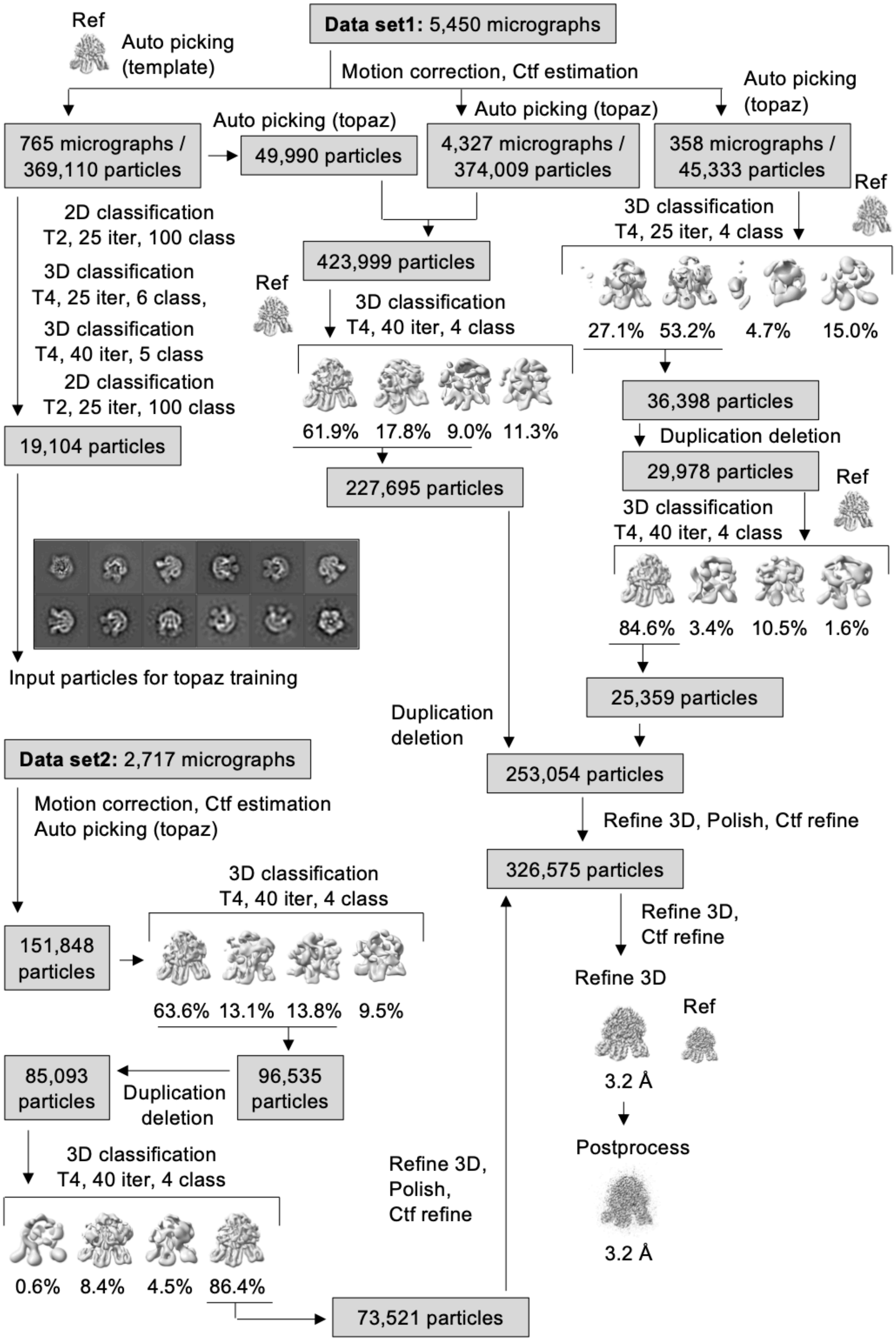


**Fig. S6. 3D reconstruction scheme of *Va*-PomAB(D24N) with NaCl.**

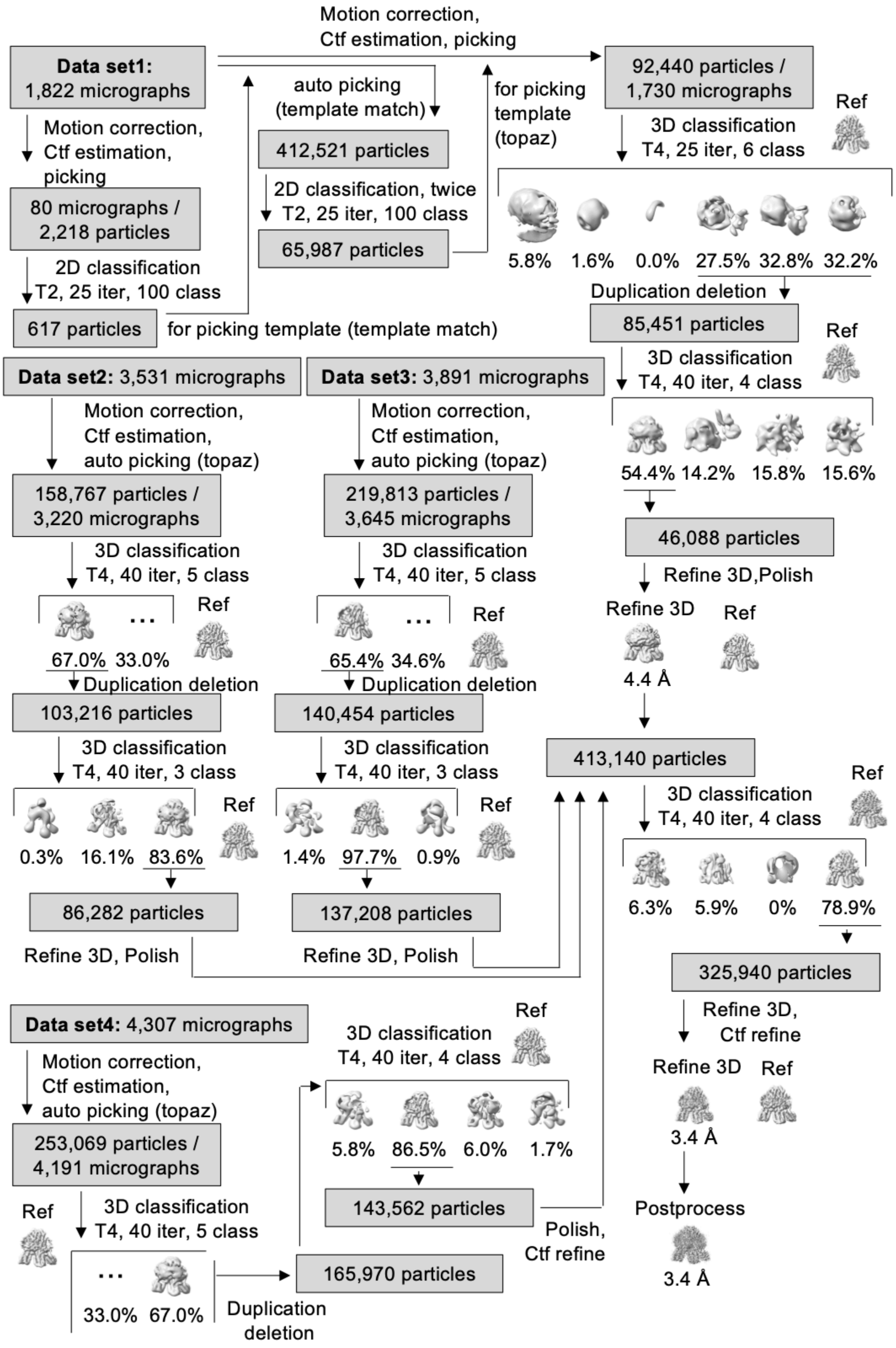


**Fig S7. 3D reconstruction scheme of *Va*-PomAB(D24N) with KCl.**

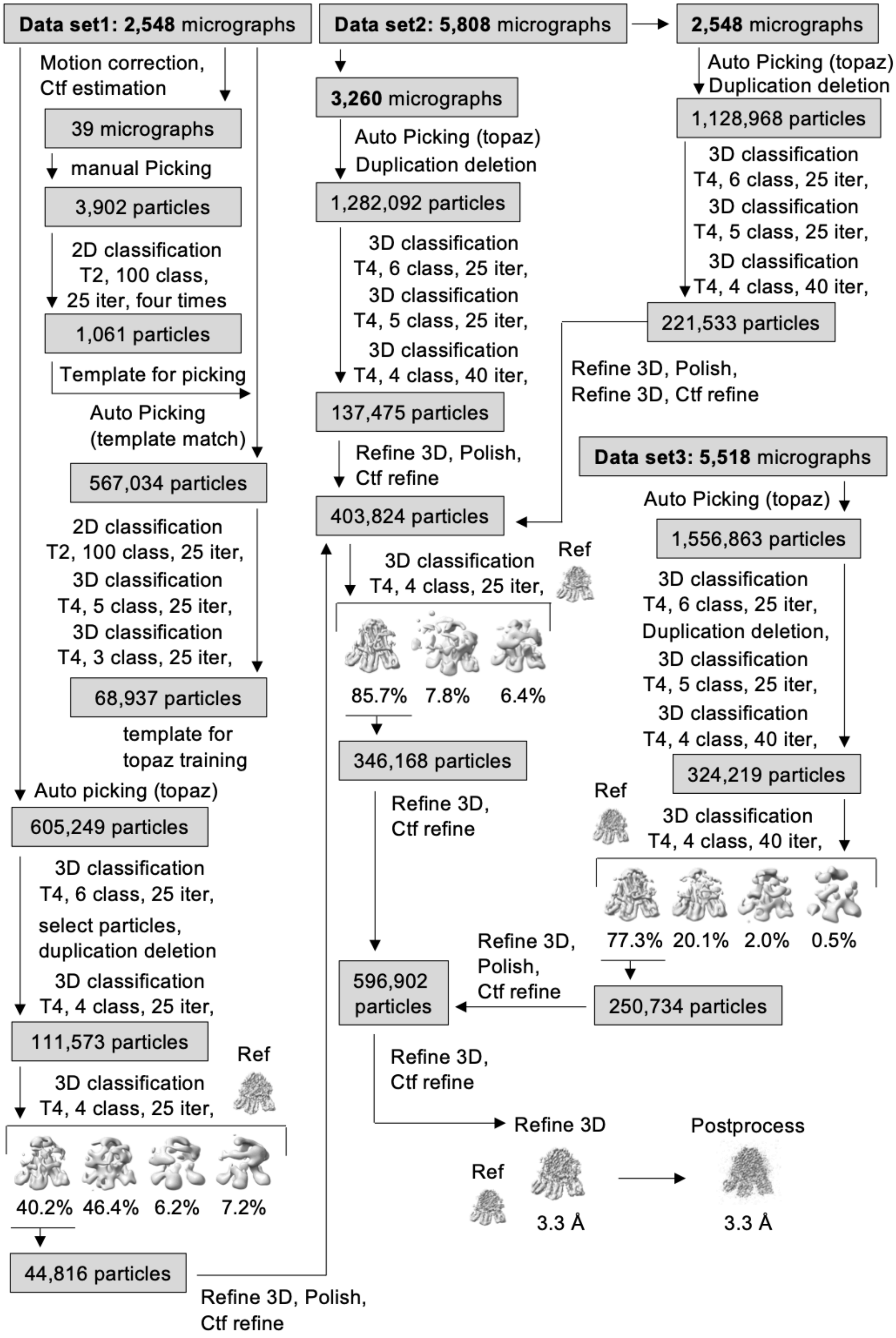


**Fig. S8. 3D reconstruction scheme of *Va*-PomAB with Phenamil.**

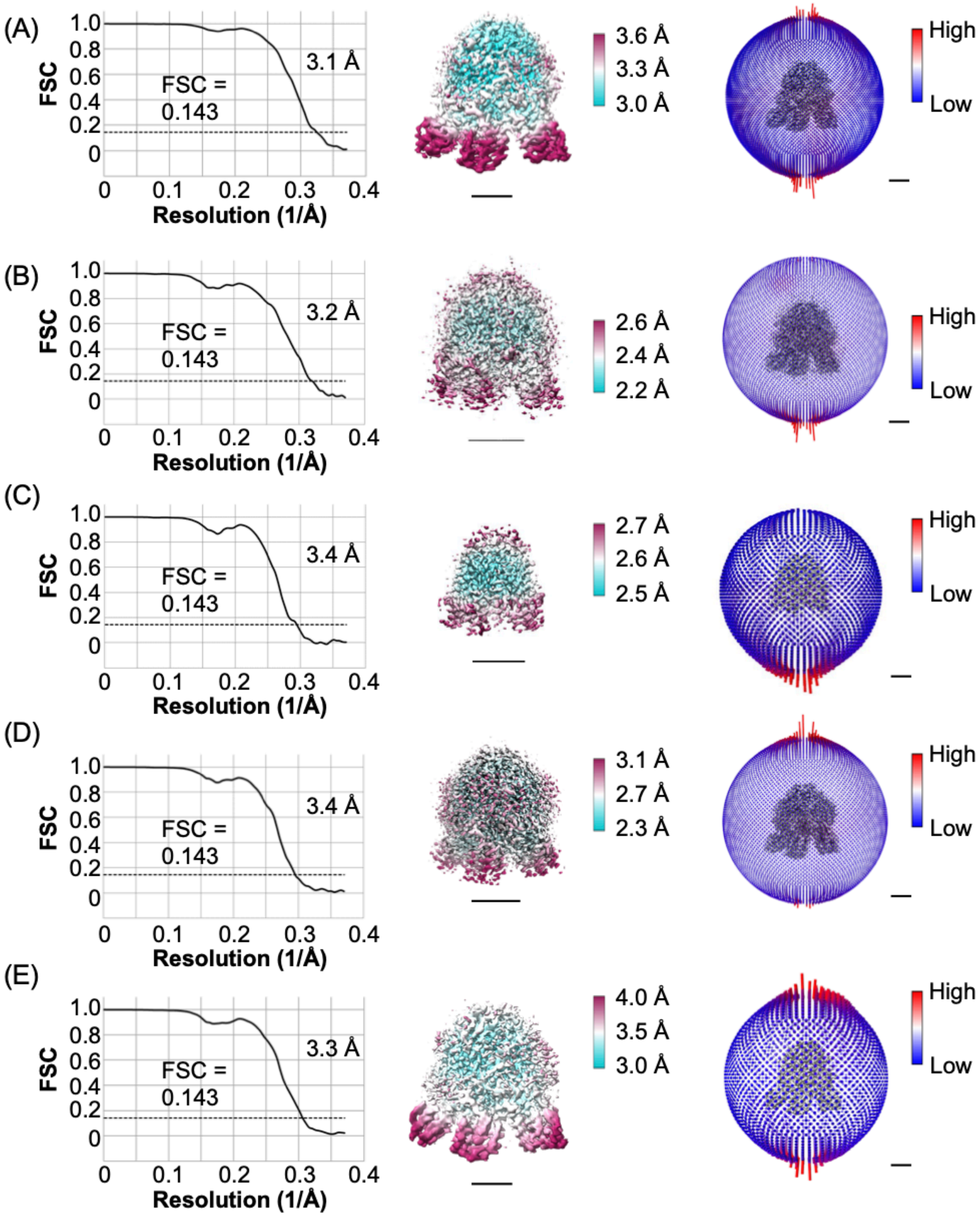


**Fig. S9. Resolution estimation of the cryo-EM map.**
The Fourier shell correlation (FSC) curve, density map colored by local resolution, and angular distribution of particles used for 3D reconstruction are shown in the left, middle, and right panels, respectively. (A) *Va*-PomAB with NaCl, (B) *Va*-PomAB(D24N) with NaCl, (C) *Va*-PomAB with KCl, (D) *Va*-PomAB(D24N) with KCl, and (E) *Va*-PomAB with Phenamil.

**Table S1. Strains and plasmids used in this study.**

| Strains or plasmids | Genotype or description | Reference or source |
|---|---|---|
| ***V. alginolyticus*** | | |
| VIO5 | Wild type strain of a polar flagellum ($Rif^r$ $Pof^+$ $Laf^-$) | (*56*) |
| NMB191 | VIO5 Δ*pomApomB* ($Rif^r$ $Laf^-$ $Mot^-$) | (*57*) |
| ***E. coli*** | | |
| DH5α | $F^-$, Φ80d*lacZ*ΔM15, Δ(*lacZYA*-*argF*)U169, *deoR*, *recA*1, *endA*1, *hsdR*17($r_K^-$, $m_K^+$), *phoA*, *supE*44, $\lambda^-$, *thi*-1, *gyrA*96, *relA*1<br>(Host for cloning experiments) | (*58*) |
| BL21(DE3)/ pLysS | $F^-$, *ompT, hsdS*$_B$($r_B^-$ $m_B^-$), *gal*( *λ cI* 857, *ind1*, *Sam7*, *nin5, lacUV*5-T7*gene*1), *dcm*(DE3)<br>(Host for protein expression) | Novagen |
| **Plasmids** | | |
| pBAD33 | $Cm^r$, $P_{BAD}$ | (*59*) |
| pHFAB | *pomA* and *pomB* in pBAD33 | (*60*) |
| pTSK37 | *pomA* and $pomB_{\Delta 41\text{-}120}$ in pBAD33 | (*17*) |
| pColdIV | Cold shock expression vector, $Amp^r$ | Takara |
| pCold4-pomAB-$His_6$ | *pomA* and *pomB-his*$_6$ in pColdIV | (*30*) |

$Rif^r$, rifampin resistant; $Pof^+$, normal polar flagellum; $Laf^-$, defective in lateral flagellar formation; $Mot^-$, defective in polar flagellar motility; $Cm^r$, chloramphenicol resistance; $P_{BAD}$, arabinose promoter; $Amp^r$, Ampicillin resistance.

**Table S2. summary of data correction and model statistics**

| **Data collection and processing** | *Va*-PomAB<br>0.1 M NaCl<br>(EMD-60580)<br>(PDB 8ZYV) | *Va*-PomAB<br>0.1 M KCl<br>(EMD-60581)<br>(PDB 8ZYW) | *Va*-PomAB<br>(D24N)<br>0.1 M NaCl<br>(EMD-60584)<br>(PDB 8ZYZ) | *Va*-PomAB<br>(D24N)<br>0.1 M KCl<br>(EMD-60585)<br>(PDB 8ZZ0) | *Va*-PomAB<br>0.1 M NaCl<br>with phenamil<br>(EMD-60636)<br>(PDB 9IJM) |
|---|---|---|---|---|---|
| EM equipment | Titan Krios | Titan Krios | Titan Krios | Titan Krios | Titan Krios |
| Detector | Gatan K3 | Gatan K3 | Gatan K3 | Gatan K3 | Gatan K3 |
| Energy filter | Gatan GIF Quantum, 20 eV slit | Gatan GIF Quantum, 20 eV slit | Gatan GIF Quantum, 20 eV slit | Gatan GIF Quantum, 20 eV slit | Gatan GIF Quantum, 20 eV slit |
| Ligand | - | - | - | - | 0.01 mM Phenamil |
| Magnification | 105,000 | 105,000 | 105,000 | 105,000 | 105,000 |
| Voltage (kV) | 300 | 300 | 300 | 300 | 300 |
| Pixel size (Å) | 0.675 | 0.675 | 0.675 | 0.675 | 0.675 |
| Symmetry imposed | C1 | C1 | C1 | C1 | C1 |
| **Data set1**<br>Micrographs (no.)<br>Electron exposure ($e^- Å^{-2}$)<br>Defocus range (μm)<br>Initial particle images (no.) | <br>1,248<br>50<br>-1.8~-0.8<br>127,822 | <br>3,947<br>60<br>-1.8~-0.6<br>2,220,744 | <br>5,450<br>60<br>-1.8~-0.6<br>788,452 | <br>1,822<br>60<br>-2.2~-0.8<br>92,440 | <br>2,548<br>60<br>-2.0~-0.725<br>605,249 |
| **Data set2**<br>Micrographs (no.)<br>Electron exposure ($e^- Å^{-2}$)<br>Defocus range (μm)<br>Initial particle images (no.) | <br>3,318<br>60<br>-2.4~-1.125<br>465,830 | <br>13,730<br>60<br>-1.8~-0.6<br>2,409,941 | <br>2,717<br>60<br>-1.8~-0.6<br>151,848 | <br>3,531<br>60<br>-2.2~-0.8<br>158,767 | <br>5,808<br>60<br>-2.8~-0.8<br>2,411,060 |
| **Data set3**<br>Micrographs (no.)<br>Electron exposure ($e^- Å^{-2}$)<br>Defocus range (μm)<br>Initial particle images (no.) | <br>5542<br>60<br>-2.4~-1.125<br>1,865742 | - | - | <br>3,891<br>60<br>-2.2~-0.8<br>219,813 | <br>5,518<br>60<br>-2.0~-0.8<br>1.556,863 |
| **Data set4**<br>Micrographs (no.)<br>Electron exposure ($e^- Å^{-2}$)<br>Defocus range (μm)<br>Initial particle images (no.) | <br>7,181<br>60<br>-1.6~-0.6<br>693,855 | - | - | <br>4,307<br>60<br>-2.0~-0.8<br>253,069 | - |

| Software | Relion 3.1 | Relion 3.1 & 4β | Relion 4β | Relion 4β | Relion 4β |
|---|---|---|---|---|---|
| Final particle images (no.) | 662,015 | 255,000 | 326,575 | 325,940 | 596,902 |
| Map Resolution (Å)<br>FSC threshold | 3.1<br>0.143 | 3.4<br>0.143 | 3.2<br>0.143 | 3.4<br>0.143 | 3.3<br>0.143 |
| **Refinement** | | | | | |
| Initial model used (PDB code) | 6YSL | 8ZYV | 8ZYV | 8ZYV | 8ZYV |
| Model resolution (Å)<br>FSC threshold | 3.1<br>0.5 | 3.4<br>0.5 | 3.2<br>0.5 | 3.4<br>0.5 | 3.3<br>0.5 |
| Map sharpening B factor (Å$^2$) | -133.426 | -131.963 | -114.517 | -135.943 | -125.242 |
| **Model composition**<br>Non-hydrogen atoms<br>Protein residues<br>Solvent<br>Ligands | <br>8,913<br>1,174<br>5<br>2 | <br>8,910<br>1,174<br>4<br>0 | <br>8,918<br>1,174<br>5<br>2 | <br>8,914<br>1,174<br>3<br>0 | <br>8,713<br>1,144<br>1<br>3 |
| **r.m.s. deviations**<br>Bond lengths (Å)<br>Bond angles (°) | <br>0.003<br>0.488 | <br>0.003<br>0.5456 | <br>0.004<br>0.529 | <br>0.003<br>0.582 | <br>0.006<br>0.689 |
| **Validation**<br>MolProbity score<br>Clashscore<br>Poor rotamers (%) | <br>1.4<br>7.29<br>0.21 | <br>1.47<br>8.67<br>0.32 | <br>1.49<br>8.66<br>0.11 | <br>1.51<br>9.76<br>0 | <br>1.68<br>12.44<br>0.44 |
| **Ramachandran plot statistics (%)**<br>Preferred<br>Allowed<br>Outlier | <br>98.4<br>1.6<br>0 | <br>98.3<br>1.7<br>0 | <br>97.9<br>2.1<br>0 | <br>98.1<br>1.9<br>0 | <br>97.7<br>2.3<br>0 |
